# The intracellular symbiont *Wolbachia* alters *Drosophila* development and metabolism to buffer against nutritional stress

**DOI:** 10.1101/2023.01.20.524972

**Authors:** Amelia RI Lindsey, Jason M Tennessen, Michael A Gelaw, Megan W Jones, Audrey J Parish, Irene LG Newton, Travis Nemkov, Angelo D’Alessandro, Madhulika Rai, Nicole Stark

## Abstract

The intracellular bacterium *Wolbachia* is a common symbiont of many arthropods and nematodes, well studied for its impacts on host reproductive biology. However, its broad success as a vertically transmitted infection cannot be attributed to manipulations of host reproduction alone. Using the *Drosophila melanogaster* model and their natively associated *Wolbachia* strain “*w*Mel”, we show that *Wolbachia* infection supports fly development and buffers against nutritional stress. *Wolbachia* infection across several fly genotypes and a range of nutrient conditions resulted in reduced pupal mortality, increased adult emergence, and larger size. We determined that the exogenous supplementation of pyrimidines partially rescued developmental phenotypes in the *Wolbachia*-free flies, and that *Wolbachia* titers were responsive to reduced gene expression of the fly’s *de novo* pyrimidine synthesis pathway. In parallel, transcriptomic and metabolomic analyses indicated that *Wolbachia* impacts larval biology far beyond pyrimidine metabolism. *Wolbachia*-infected larvae had strong signatures of shifts in glutathione and mitochondrial metabolism, plus significant changes in the expression of key developmental regulators including *Notch*, the insulin receptor (*lnR*), and the juvenile hormone receptor *Methoprene-tolerant* (*Met*). We propose that *Wolbachia* acts as a beneficial symbiont to support fly development and enhance host fitness, especially during periods of nutrient stress.

**SIGNIFICANCE:** *Wolbachia* is a bacterial symbiont of arthropods and nematodes, well described for its manipulations of arthropod reproduction. However, many have theorized there must be more to this symbiosis, even in well-studied *Wolbachia-*host relationships such as with *Drosophila*. Reproductive impacts alone cannot explain the success and ubiquity of this bacterium. Here, we use *Drosophila melanogaster* and their native *Wolbachia* infections to show that *Wolbachia* supports fly development and significantly buffers flies against nutritional stress. These developmental advantages might help explain the ubiquity of *Wolbachia* infections.

## INTRODUCTION

Maternally transmitted microbes have evolved numerous ways to alter host biology, ultimately facilitating their own transmission. Strategies include supplementation of the host’s diet via nutrient provisioning, protection against parasites and infections, and even direct manipulation of host reproduction (1). Many insects, other terrestrial arthropods, and some nematode species are infected with bacteria in the genus *Wolbachia* (Alphaproteobacteria: Rickettsiales), a maternally transmitted infection long appreciated for its direct impacts on host reproduction and sex ratios (1–10). Critically, *Wolbachia* can spread through populations in the absence of such reproductive manipulations (11). And, increasingly, other *Wolbachia*-mediated benefits to the host are being uncovered (12). For example, a suite of *Wolbachia* strains can protect host insects against secondary infections, especially viruses, a phenomenon that now forms the basis of many ongoing vector control programs across the world (13, 14). Certain strains of *Wolbachia* are obligate beneficial nutritional symbionts, such as the bed bug-infecting *Wolbachia* that produce B-vitamins to support their obligately hematophagous hosts (15). In other insects, facultative associations with *Wolbachia* have likewise been implicated in the balance of cholesterol, iron, or B-vitamins (16–19).

We previously identified that *Wolbachia* infection results in a suite of changes to the expression of *Drosophila melanogaster* nucleotide metabolism pathways (20). Additionally, comparative genomic data indicate that the biosynthetic pathways for the *de novo* synthesis of purines and pyrimidines have been conserved across *Wolbachia*, pointing towards their broad importance (21). Furthermore, *Wolbachia* encode for an abundance of amino acid importers, which would provide necessary precursors for *Wolbachia*’s *de novo* synthesis of nucleotides (3, 21). The conservation of these metabolic pathways might simply represent essential functions for *Wolbachia*’s own metabolism, or they might also be important in the context of interaction with the host.

While there are increasing examples in the literature that hint at significant metabolic interplay between *Wolbachia* and hosts, our understanding of the symbiosis is still largely focused on reproductive processes and other aspects of adult biology (1). Indeed, we know very little about *Wolbachia*’s interactions with hosts during development. In insects, not only is the juvenile period physiologically distinct (especially in holometabolous insects that undergo complete metamorphosis) but also this developmental period determines many aspects of adult biology and fitness (22–24). To address this significant gap in knowledge, we undertook a series of experiments using the fruit fly, *Drosophila melanogaster*, naturally infected with the *Wolbachia* strain *w*Mel. We hypothesized that *Wolbachia* plays a supporting role in host nutrition, and used a combination of transcriptomics, metabolomics, diet manipulations, and fly genetics to investigate the relationship between developing flies and their *Wolbachia* infections.

## METHODS

### Fly husbandry

*Drosophila melanogaster* stocks with or without their native *Wolbachia* strain “*w*Mel” were maintained on standard Bloomington cornmeal-agar medium at 25 °C on a 24-hour, 12:12 light:dark cycle under density-controlled conditions and 50% relative humidity. *Wolbachia* colonization status was confirmed with PCR assays using *Wolbachia*-specific 16S primers WspecF and WspecR (25). Genotypes used in nutritional assays (below) included: DGRP-320 (RRID:BDSC_29654), a *Wolbachia*-infected isogenic wild-type strain with genome sequence available which we refer to as “wild-type” below (26), and a *Wolbachia*-infected white line, *w*^145^ (RRID:BDSC_145). *Wolbachia*-cleared counterpart stocks were generated with antibiotics via three generations of tetracycline treatment (20 μg/mL in the fly food for three generations), followed by re-inoculation of the gut microbiome by transfer to bottles that previously harbored male flies from the original stock that had fed and defecated on the media for one week (27). Stocks were allowed to recover from any potential transgenerational effects of the antibiotic treatment for at least an additional ten generations prior to use in any experiment and re-screened for *Wolbachia* infection immediately prior to all experiments. All other fly stocks (e.g., RNAi lines), are detailed in Supplementary Table S1.

### RNA-Seq of larvae

Pools of second instar (“L2”) DGRP-320 larvae with or without *Wolbachia* were processed for transcriptomic analyses. For each infection status, three vials were initiated by allowing five pairs of adult flies per vial to mate and lay eggs on standard Bloomington cornmeal-agar medium for 8 hours. After this period, the adults were removed, and larvae were maintained under standard rearing conditions. Flies were allowed to develop until the L2 stage, at which point they were removed from each vial with a dissecting pin and transferred to a microcentrifuge tube containing sterile 1X phosphate buffered saline (PBS). Each pool of 20 L2s was washed three times with 1X PBS, after which the solution was removed, and larvae were flash frozen in liquid nitrogen. Total RNA was extracted from each pool using the Monarch® Total RNA Miniprep Kit (New England Biolabs) including on-column DNase treatment. RNA-seq libraries were prepared by Novogene Co, Ltd. and followed a standard strand-specific Illumina-compatible preparation (New England Biolabs) including purification of mRNA using oligo-dT beads. Libraries were sequenced on an Illumina NovaSeq platform to generate ∼9Gb of paired end 150bp reads per library. Reads were mapped to extracted reference transcripts of the *Drosophila melanogaster* reference genome (release 6.48) (28) using the RSEM v. 1.3.0 (29) programs ‘rsem-prepare-reference’ and ‘rsem-calculate-expression’, employing the STAR v. 2.3.5a aligner (30). Transcript abundance was summarized and imported to R v. 4.3.0 (31) with tximport v. 1.28.0 (32) for use in downstream analyses. Only genes with >1 count per million in at least two samples were retained for analysis. Differential gene expression was assessed with EdgeR v. 3.42.4 (33, 34), employing a TMM normalization, dispersion calculation, and a generalized linear model, with quasi-likelihood F-tests (function ‘glmQLFit’). Genes that were significantly differentially expressed were defined as those with a false discovery rate (FDR) q-value of <0.05. Mitochondrial genes in the DEG dataset were extracted along with their annotations in mitoXplorer 2.0 (35, 36). We determined enrichment with PANGEA (37) using all DEGs as the test set, all expressed genes (Supplemental Table S3) as the background, and gene set categories of ‘KEGG Pathway D. mel’, ‘Preferred tissue (modEncode RNA_seq)’, and all three *Drosophila* GOslim gene sets (Biological Process, Cellular Component, Molecular Function). Gene sets with p<0.05 after Benjamini & Hochberg corrections for multiple testing, or >2 fold-change with uncorrected p-value of p<0.05 are displayed. DEGs were also clustered into an interaction network using STRING v.1.4.2 (38), implemented in Cytoscape v.3.6.0 (39), which simultaneously assesses overrepresentation of STRING clusters. The confidence threshold for network building was set to 0.85 (i.e., highly stringent).

### *Wolbachia* developmental transcriptomics

To assess *Wolbachia* gene expression, we analyzed the *Wolbachia* transcriptomes extracted from the MODENCODE dataset (40, 41). This dataset has previously been used to find *Wolbachia* genes that are significantly differentially expressed across fly development (40–42). Importantly, and as those studies have noted, this is a large data set with limited power to detect differences due to low replication across a very large number of stages that are not necessarily biologically independent or strictly linearly correlated. As such, previously identified DEGs are primarily those with large changes in expression and limited developmental stage-by-expression interactive effects. To combat this challenge, we opted to take a genome-wide approach to look at these data without the goal of identifying specific differentially expressed genes. The dataset was further curated by removing tissue-specific libraries (e.g., L3 midguts), and averaging TPM values within a developmental time point for those represented by multiple libraries. To assign orthology to each *w*Mel protein, we used a previously constructed database (43) of proteins from 33 *Wolbachia* genomes (representing nine Supergroups: A, B, C, D, E, F, J, L, T), clustered into orthologous groups with ProteinOrtho v5.15 (44). We assessed each *w*Mel protein for phylogenetic conservation as either a “core” or “unique” gene at a range of taxonomic levels. The “unique” designation was for proteins that were found in all members of a given clade and also restricted to that clade (i.e. not found in any other *Wolbachia* strains). The “core” designation was for proteins that were present in all members of a clade without regard to their presence in other strains outside of that clade. Using these assignments we designated the following groups of *w*Mel proteins: (1) unique to *w*Mel, (2) core to the *Drosophila*-infecting clade within Supergroup A (*w*Mel, *w*Ri*, w*Au, *w*Rec, *w*Ha, *w*Suz), (3) core to Supergroup A, (4) core to Supergroups A and B, (5) core proteins across all *Wolbachia*, and, (6) all other *w*Mel proteins. Statistics and data visualization were carried out in R version 4.1.3 (31). Expression clustering and visualization was performed with R package ‘pheatmap’ (45) and clustering method ‘complete’. Functional enrichment tests for *w*Mel gene sets were performed with ShinyGO 0.80 (46) using ‘Wolbachia endosymbiont of Drosophila melanogaster’ (ID = ‘STRING.163164.Wolbachia’) as the species, a custom background gene set of the 1191 expressed genes, GO Biological Process as the pathway database, and an FDR cutoff of p = 0.05.

### Nutritional stress experiments

To generate poor media, Nutri-fly® Bloomington Formulation (Genesee Scientific 66-121) was prepared according to manufacturer’s instructions and diluted to indicated percentages while maintaining the full concentrations of agar (5.3 g/L) and propionic acid (4.8 ml/L). Five ml of media was aliquoted into each vial. Flies were placed in mating cages on grape agar plates streaked with yeast paste and acclimated for 24 hours, prior to initiating 4-hour egg lays. Upon hatching, < 4-hour old L1 larvae were transferred to media vials (n=20 per vial), and development was scored every 24 hours. Pupation was defined by eversion of the anterior spiracles (47). Each vial was tracked until all flies eclosed, or until three consecutive days without additional adult emergence.

### Axenic assays

To test if the gut microbiome was playing a role in *Wolbachia-*mediated developmental phenotypes, we reared wild-type flies with and without *Wolbachia* on 100% and 25% strength media under axenic conditions (i.e., without the gut or food microbiome). 10 mL of media (prepared according to protocols in “nutritional stress experiments”) was aliquoted into polypropylene wide vials, capped with a Cellulose Acetate Flug® (Flystuff 49-101), and autoclaved on liquid setting for 20 minutes to sterilize. Embryos were collected on grape agar with yeast paste as described previously and transferred into 70-micron mesh cell strainers (Falcon 352350). Embryos in the cell strainers were washed with *Drosophila* embryo wash solution (7% NaCl, 0.5% Triton X-100) to remove food debris. After an initial wash, embryos were immersed in a 10% bleach solution for 3 minutes, with gentle mixing every 30 seconds. The embryos were then washed with sterile *Drosophila* embryo wash solution to remove excess bleach. Embryos were subjected to a final rinse with sterile phosphate buffered saline (PBS) and transferred to a sterile agar culture plate (2% agar in deionized water) stained with blue gel food coloring to facilitate counting. A flame-sterilized probe was used to transfer 20 embryos into each vial of sterile media, and developmental assays were carried out as described above.

### Nutritional supplementation experiments

To test the impact of specific nutrients, we prepared 12.5% strength media as above, and selectively added in protein or nucleotides: the precursors and end points of *de novo* nucleotide biosynthesis. Treatments included casein (for bulk addition of amino acids at 64.1 g/L; Sigma C7078), inosine (0.74 g/L; Sigma I4125), uridine (0.67 g/L; Sigma U3003), or both inosine and uridine at the aforementioned concentrations. The concentrations of additives were calculated as 87.5% of a previously defined holidic diet (48), to adjust for the 12.5% BDSC media strength. Developmental assays were carried out as described above.

### Pupal size measurements

After selected developmental assays, pupal casings and dead pupae were removed from vials using a wet paint brush and transferred to glass slides. Pupae were imaged with brightfield microscopy on an ECHO Revolve at 4X. Images were manually annotated in Echo Labs Pro software to measure sizes. Pupal length was defined as the distance between the base of the posterior spiracles to the midway point between the anterior spiracles. Width measurements were taken at the widest part of the pupa. The volume of each pupa was calculated assuming a prolate spheroid shape [V = (4/3) π (width/2)^2^ (length/2)] (Supplemental Figure S2) (49, 50). For assays in which we sexed and measured pupae, we allowed five pairs of flies to mate and lay eggs on the appropriate media for four days, after which the adults were cleared, and offspring were allowed to develop under standard rearing conditions. Approximately 50 pupae per condition were collected at pupal stages P13-14 (51), and sexed based on the presence of sex combs prior to measurement.

### Knockdown of fly nucleotide metabolism genes

*Wolbachia*-infected Transgenic RNAi Project (“TRiP”) stocks were obtained from the Bloomington *Drosophila* Stock Center to knock down expression of genes in the purine and pyrimidine nucleotide biosynthesis pathways (Supplemental File S1). Unmated females carrying a UAS-gene-specific short hairpin were crossed to males with an Act5C-Gal4 driver (RRID:BDSC_3954: y^1^w*; P{w^+mC^=Act5C-GAL4}17bFO1/TM6B, Tb^1^). Unmated, F1 female progeny were sorted as adults into those with the short hairpin and Gal4-driver or those with the short hairpin and the TM6B balancer. At three days old, the sorted flies were flash frozen in liquid nitrogen and then stored at −80 °C for later processing. To determine the effect of the TM6B balancer on *Wolbachia* titer, we crossed *Wolbachia*-infected, Act5C-Gal4 driver females to *Wolbachia*-uninfected UAS-anti-Ppyr\LUC males (RRID:BDSC_31603: y^1^ v^1^; P{y^+t7.7^ v^+t1.8^=TRiP.JF01355}attP2). Unmated, F1 female progeny were sorted and stored as described for the nucleotide biosynthesis gene knockdowns. To determine spatial differences in *Wolbachia* titer, we dissected ovaries from two-day old *r-l* knockdown and sibling control flies. We separated ovaries from living, anesthetized flies in sterile PBS. Ovaries and carcasses from three flies were pooled for each biological replicate and flash frozen, followed by storage at −80 °C for later processing.

### Real-time quantitative RT-PCR analyses of target gene expression

Flies were homogenized in TRIzol reagent (Invitrogen), and total RNA was extracted following manufacturer’s instructions. RNA extractions were DNase treated (RQ1 RNase-free DNase, New England Biolabs) according to the manufacturer’s instructions. cDNA was synthesized using M-MuLV Reverse Transcriptase (New England Biolabs) with random hexamer primers (Integrated DNA Technologies). Quantitative RT-PCR reactions were performed with SensiFAST SYBR Hi-ROX kit (Bioline) and gene-specific primers (Supplemental File S2). All samples were run in technical duplicate alongside negative controls on an Applied Bioscience StepOnePlus qPCR machine (Life Technologies). Gene expression was normalized to endogenous 18S rRNA expression using the Livak method (52).

### Real-time quantitative PCR analyses of *Wolbachia* titer

DNA was extracted from individual flies using the Quick-DNA/RNA Pathogen Miniprep kit (Zymo Research) according to the manufacturer’s protocol. *ftsZ* primers were used to quantify *Wolbachia* genome copy numbers, which were normalized to host genome copies via 18S quantification using the Livak method (52). Reactions were performed with SensiFAST SYBR Hi-ROX kit (Bioline), and all samples and negative controls were run in technical duplicate on an Applied Bioscience StepOnePlus qPCR machine (Life Technologies).

### Metabolomics

We reared L2 larvae of both wild-type (DGRP-320) and *w*^145^ flies, with and without *Wolbachia*, on 100% and 25% BDSC media for metabolomics analyses. Larvae were collected and processed for metabolomics following published protocols and best practices (53, 54). For each of the eight conditions (genotype\**Wolbachia*\*media), a minimum of six vials were initiated by allowing five pairs of adult flies in each vial to mate and lay eggs for 8 hours. After this period, the adults were removed, and larvae were maintained under standard rearing conditions. Flies were allowed to develop until the L2 stage, at which larvae were stage-verified, picked from the media with a dissecting pin, and transferred to a microcentrifuge tube containing sterile ice-cold 0.9% sodium chloride (NaCl). Approximately 30 larvae originating from the same vial of media were collected and pooled into a single replicate, resulting in a minimum of six independent biological replicates per condition. Larvae were rinsed in 0.9% ice cold NaCl three times, gently pelleted, the liquid was removed, and then the pool of larvae was immediately flash frozen in liquid nitrogen prior to storing at −80 °C for later processing. All pools were processed in under 20 minutes to limit decay of metabolites. For metabolite extraction, samples were first transferred to tared 2 mL screwcap tubes containing 1.4 mm ceramic beads pre-chilled in liquid nitrogen. The sample mass was recorded, and tubes were immediately placed back in liquid nitrogen. 800 mL of prechilled (−20 °C) 90% methanol containing 2 µg/mL succinic-d4 acid was added to each sample tube and the sample was homogenized in an Omni Beadruptor 24 for 30 seconds at 6.4 m/s. The samples were removed from the homogenizer, incubated at −20°C for 1 hr, and centrifuged at 20,000 x g for 5 min at 4 °C. 600 μl of the supernatant was transferred into a new 1.5 mL microcentrifuge tube and dried overnight in a vacuum centrifuge. Ultra High-Pressure Liquid Chromatography -Mass Spectrometry (UHPLC-MS)-based Metabolomics analyses were performed at the University of Colorado Anschutz Medical Campus, as previously described (55). Briefly, the analytical platform employs a Vanquish UHPLC system (Thermo Fisher Scientific, San Jose, CA, USA) coupled online to a Q Exactive mass spectrometer (Thermo Fisher Scientific, San Jose, CA, USA). The (semi)polar extracts were resolved over a Kinetex C18 column, 2.1 x 150 mm, 1.7 µm particle size (Phenomenex, Torrance, CA, USA) equipped with a guard column (SecurityGuard^TM^ Ultracartridge – UHPLC C18 for 2.1 mm ID Columns – AJO-8782 – Phenomenex, Torrance, CA, USA) using an aqueous phase (A) of water and 0.1% formic acid and a mobile phase (B) of acetonitrile and 0.1% formic acid for positive ion polarity mode, and an aqueous phase (A) of water:acetonitrile (95:5) with 1 mM ammonium acetate and a mobile phase (B) of acetonitrile:water (95:5) with 1 mM ammonium acetate for negative ion polarity mode. The Q Exactive mass spectrometer (Thermo Fisher Scientific, San Jose, CA, USA) was operated independently in positive or negative ion mode, scanning in Full MS mode (2 μscans) from 60 to 900 m/z at 70,000 resolution, with 4 kV spray voltage, 45 sheath gas, 15 auxiliary gas. Calibration was performed prior to analysis using the Pierce^TM^ Positive and Negative Ion Calibration Solutions (Thermo Fisher Scientific). Acquired data was converted from raw to mzXML file format using Mass Matrix (Cleveland, OH, USA). Samples were analyzed in randomized order with a technical mixture injected after every 24 samples to qualify instrument performance. Metabolite assignments were performed using accurate intact mass (sub-10 ppm), isotopologue distributions, and retention time/spectral comparison to an in-house standard compound library (MSMLS, IROA Technologies, NJ, USA) using El-MAVEN (Elucidata, San Francisco, CA, USA).

### Statistics and Data Visualization

Statistics and data visualization were carried out in R version 3.5.0 (31). Significant differences in development were assessed with generalized linear mixed-effects models (package: ‘lme4’, function ‘glmer’ (56)) including the proportion of flies that reached a given stage (pupa or adult) as a binomial response, *Wolbachia* presence, day of development, media, and the interaction of the three as fixed effects, and vial as a random effect to account for repeated measures. Significant differences in pupal volume of flies reared on 12.5% media was assessed with a two-way ANOVA (function ‘aov’) including *Wolbachia*, mortality status, and their interaction as fixed effects. Significant differences in sizes of pupae reared on supplemented media was first assessed with a two-way ANOVA (function ‘aov’) including *Wolbachia*, media, and their interaction as fixed effects. Paired comparisons between *Wolbachia*-infected and uninfected flies were then assessed with Wilcoxon rank sum tests (function ‘wilcox.test’). Gene expression and *Wolbachia* titers from knockdown experiments were assessed with two-way ANOVAs (function ‘aov’) including delta delta Ct as the response, and target locus, genotype (knockdown versus sibling), and their interaction as fixed effects. *Wolbachia* titers in dissected flies were also assessed with ANOVA, here with genotype, tissue, and their interaction as fixed effects. Pairwise comparisons were performed with t-tests. Similarity of metabolomics samples were visualized with non-metric multidimensional scaling (NMDS) using the ‘metaMDS’ function from the R ‘vegan’ package (57) and euclidean distances. Sample collection and data processing for the DGRP-320 and *w*^145^ flies were carried out on separate dates so genotypes were analyzed separately. Differential abundance of metabolites was determined with MetaboAnalyst 6.0 (58) and leveraged normalization by sample mass, square root transformation, and pareto scaling followed by default analysis parameters.

## RESULTS

### *Wolbachia* infection drastically alters larval gene expression

Given that the *Wolbachia*-host relationship is relatively unexplored in juvenile insects, we used a transcriptomics approach to broadly investigate the differences in larval biology when flies are infected with *Wolbachia*. At the second instar (L2) stage, 1,477 genes (∼14% of all expressed genes) were significantly differentially expressed (“DEGs”) between the *Wolbachia* infected and uninfected flies (786 upregulated; 691 downregulated in the presence of *Wolbachia*; see Supplemental Table S4). DEGs had as much as ∼100-fold change in expression. Included in the set of DEGs were two critical regulators of development, both downregulated in the presence of *Wolbachia*: *notch* and the insulin receptor *lnR.* Additional related DEGs included *lst* (a peptide hormone that suppresses insulin production, upregulated), *apolipophorin* (downregulated), *smr* (mediates repression of ecdysone and Notch signaling, downregulated), *ecdysis triggering hormone* and its receptor (*eth* and *ethr*, both upregulated), and the juvenile hormone (JH) receptor *met* (upregulated), plus a suite of JH binding proteins (*jhbp4*, −11, y12, and −16, all upregulated except for *jhbp11*).

Many proteins involved in mitochondrial biology were also significantly differentially expressed (Figure 1). These included several mitochondrial ATP synthases (all downregulated), mitochondrial replication, transcription, and translation related proteins (e.g., mtDNA-helicase, MTPAP, mEFG1, mRpS10, LeuRS-m; all upregulated), heat shock proteins, transporters, and a suite of other enzymes involved in various aspects of mitochondrial metabolism (e.g., NADH dehydrogenase ND-20L, AIF, Gpo1, Sardh, mt:ND6, Acly, Taz, Tpi, amongst others). Importantly, the DEGs included several genes encoding components of the electron transport chain (Cyt-c-d, COX4L, COX5BL, COX7AL; all between 9-to 12-fold downregulated in *Wolbachia* infected larvae). Perhaps functionally related to the impacts on mitochondrial physiology, glutathione metabolism was largely upregulated in the *Wolbachia* infected flies (e.g., *gss1*, *gss2*, *gstE6*, *gstE9*, *gclm*, *gclc*, *gclm*), consistent with a number of previous reports that *Wolbachia* alters redox balance (13).

**Figure 1.**
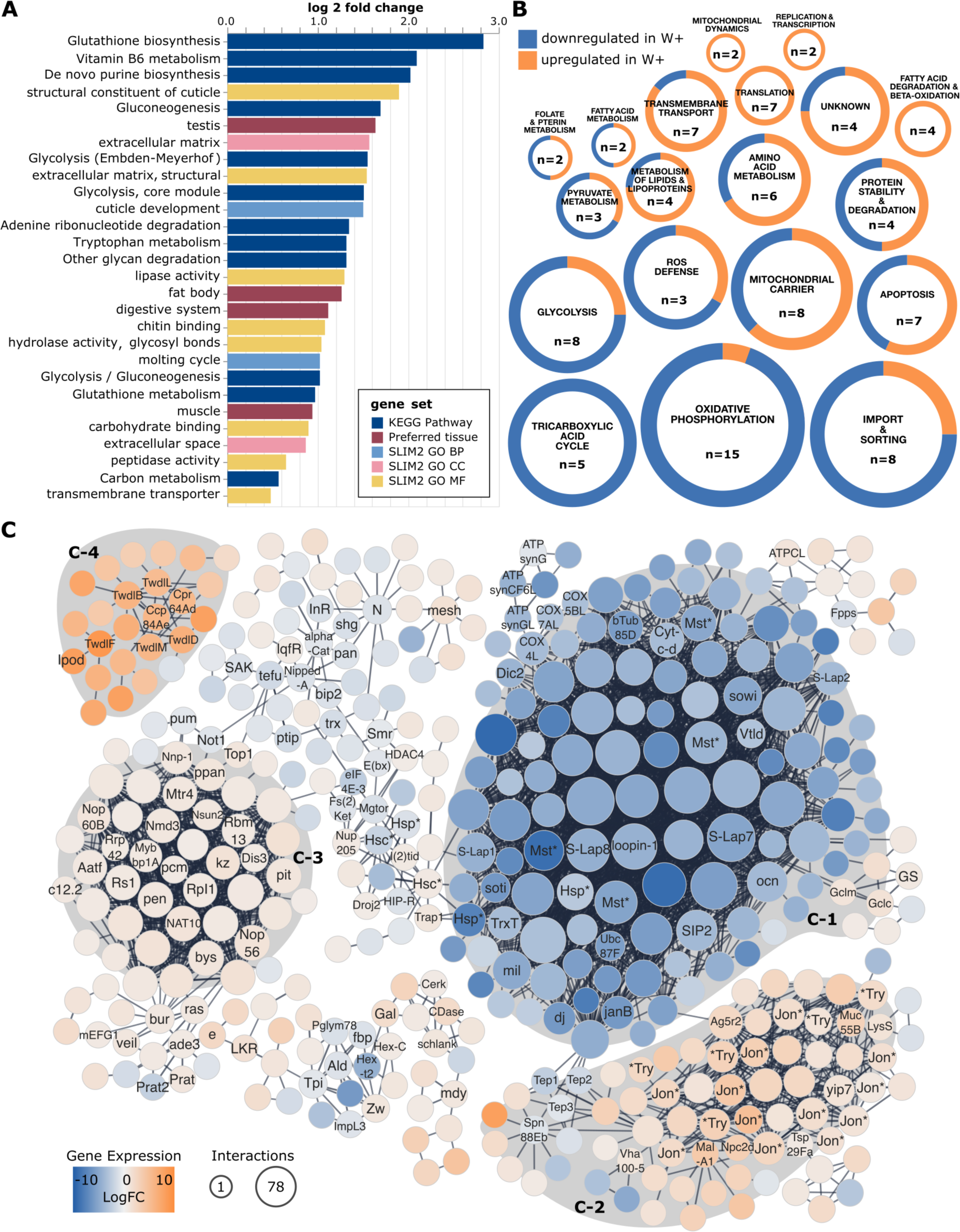
*Wolbachia* infection is associated with differential expression of 14% of the larval transcriptome. **(A)** Significantly enriched gene sets within the DEGs were assessed for enrichment using PANGEA (37). Log2 fold change values indicate the amount of enrichment for a given gene set, and color coding corresponds to the database of origin for a given gene set. Gene Ontology (GO) gene sets are the *Drosophila* GO Subsets (GO slim) for Biological Process (BP), Cellular Component (CC), and Molecular Function (MF). **(B)** Mitochondrial related DEGs. Each circle represents a functional group, as defined by mitoXplorer, with colored slices indicating the proportion of genes in that group that were up- or down-regulated in response to *Wolbachia* infection. The values “n=” in each circle indicate the number of DEGs in that group, and the size of the circle is a relative indicator of the level of overrepresentation as compared to genomic background. **(C)** Core protein-protein interaction networks within the differentially expressed gene set. Nodes (proteins) are colored according to their change in gene expression relative to uninfected larvae. The size of each node corresponds to the number of interactions that protein has with others in the network. Nodes are labeled if the gene has a name, and either (1) has greater than two connections, or, (2) if the gene was discussed elsewhere in the main text. Some gene names are abbreviated using an asterisk to denote variations on the base family gene name. These include Jon* (e.g., Jon99Fii, Jon66Ci, etc), Mst*, Hsp*, Hsc*, *Try (e.g., alphaTry, lambdaTry, etc). The full network with all nodes annotated with their full gene names is available as Supplemental Figure S1. Shaded regions denote clusters of the gene expression network that are significantly overrepresented with the following functional terms: **(C-1)** uncharacterized peptidases, transmembrane transport, protein localization to microtubule, **(C-2)** signal, serine proteases, extracellular, hydrolase, digestion, carboxypeptidase, transmembrane transport, **(C-3)** ribosome biogenesis and tRNA modification, and **(C-4)** signal, cuticle development, chitin. Full details of all categories and statistical tests for the data represented in panels A and B are in Supplemental Tables S5 and 6, respectively.

Notably, we also identified significant impacts of *Wolbachia* infection on the Toll and IMD immune pathways in the presence of *Wolbachia*. This includes downregulation of the Toll activator *spatzle* and its processing enzyme (SPE); downregulation of antimicrobial proteins including *attacins* (A, B, and D), *defensin*, and *baramicin-A1*; and differential expression of two pattern recognition receptors: peptidoglycan recognition proteins *pgrp-sa* (downregulated) and *pgrp-lb* (upregulated). We also saw a significant overrepresentation of unnamed genes: 36% of all the expressed genes were unnamed (e.g., only have the CG designation), whereas 47% of the DEGs were unnamed (X-squared = 69.12, df = 1, p < 0.0001).

To provide a more holistic look at the list of DEGs, we used two approaches: (1) over-representation analyses, and (2) network visualization based on protein-protein interactions. The DEGs were significantly overrepresented for 29 different gene sets including 13 KEGG pathways, 8 molecular functions, 4 preferred tissues, 2 biological processes (cuticle and molting), and 2 cellular components (both extracellular) (Supplemental Table S5, Figure 1A). Many overrepresented KEGG pathways have connections to mitochondrial biology and carbohydrate metabolism (e.g., glutathione biosynthesis/metabolism, glycolysis, gluconeogenesis, carbon metabolism, glycan degradation). Additionally, overrepresented KEGG pathways include vitamin B6 metabolism, de novo purine biosynthesis, adenine degradation, and tryptophan metabolism. Other overrepresented gene sets largely point towards digestion and nutrient storage. For example, preferred tissues included the digestive system, fat body (the primary site of fat storage), and muscle (the primary site of carbohydrate storage), and many overrepresented molecular functions were enzymatic (e.g., hydrolases, peptidases, lipases).

Given these signatures of altered mitochondrial biology, we specifically extracted mitochondrial processes from the DEG set and assessed expression across different functions (Figure 1B). A wide range of mitochondrial processes were represented in the DEG set. The most abundant DEGs, both in absolute number and degree of over-representation, were those related to oxidative phosphorylation: largely downregulated. Energy generation processes such as glycolysis and the tricarboxylic acid (TCA) cycle were similarly overrepresented and also largely downregulated. In contrast, transport-related functions (e.g., transmembrane transport and carriers) and core mitochondrial processes such as translation, transcription and replication, amino acid and protein metabolism, and mitochondrial dynamics were primarily upregulated. These data point to two contrasting impacts of *Wolbachia* infection on mitochondrial function: an up regulation of core processes but a down regulation of carbon and energy metabolism.

Network analyses revealed that 418 of the DEGs (28.3%) form two high confidence networks (Figure 1C, Supplemental Figure S1), each of which was also associated with significantly overrepresented STRING clusters (Supplemental Table S6). Because the list is long and somewhat redundant due to the nested nature of functional annotations, we summarize the findings as they relate to Figure 1C below. The larger network contains three clusters, each with proteins corresponding to different significantly overrepresented functional categories (Figure 1C, 1-3). The “C-1” cluster is strongly downregulated in *Wolbachia*-infected flies and contains many of the aforementioned unnamed genes. While unnamed, many of these proteins are predicted peptidases. DEGs were significantly overrepresented for terms related to transmembrane transport and protein localization to microtubule, and the related genes are also largely clustered in “C-1”. The “C-2” cluster is primarily upregulated in the presence of *Wolbachia*. Significantly overrepresented terms that cluster in C-2 include signal, serine proteases (e.g. the Jonah proteases, “Jon*”), extracellular, hydrolase, digestion, carboxypeptidase, and transmembrane transport *). The “C-3” cluster is also upregulated in the presence of *Wolbachia*. Significantly overrepresented terms here are related to translation, specifically ribosome biogenesis and tRNA modification. While not significantly overrepresented, there were many DEGs related to purine nucleotide metabolism: these can be seen below cluster C-3 and include genes such as *prat*, *prat2*, and *ade3*. Finally, the second, smaller network (Figure 2C-4) is strongly upregulated in the presence of *Wolbachia* and made up of proteins that correspond to the significantly overrepresented terms signal, cuticle development, and chitin. Interestingly, this network also includes *ipod*, the interacting <u>p</u>artner <u>o</u>f <u>D</u>nmt2 (an RNA methyltransferase previously implicated in *Wolbachia*-mediated pathogen blocking (59)).

**Figure 2.**
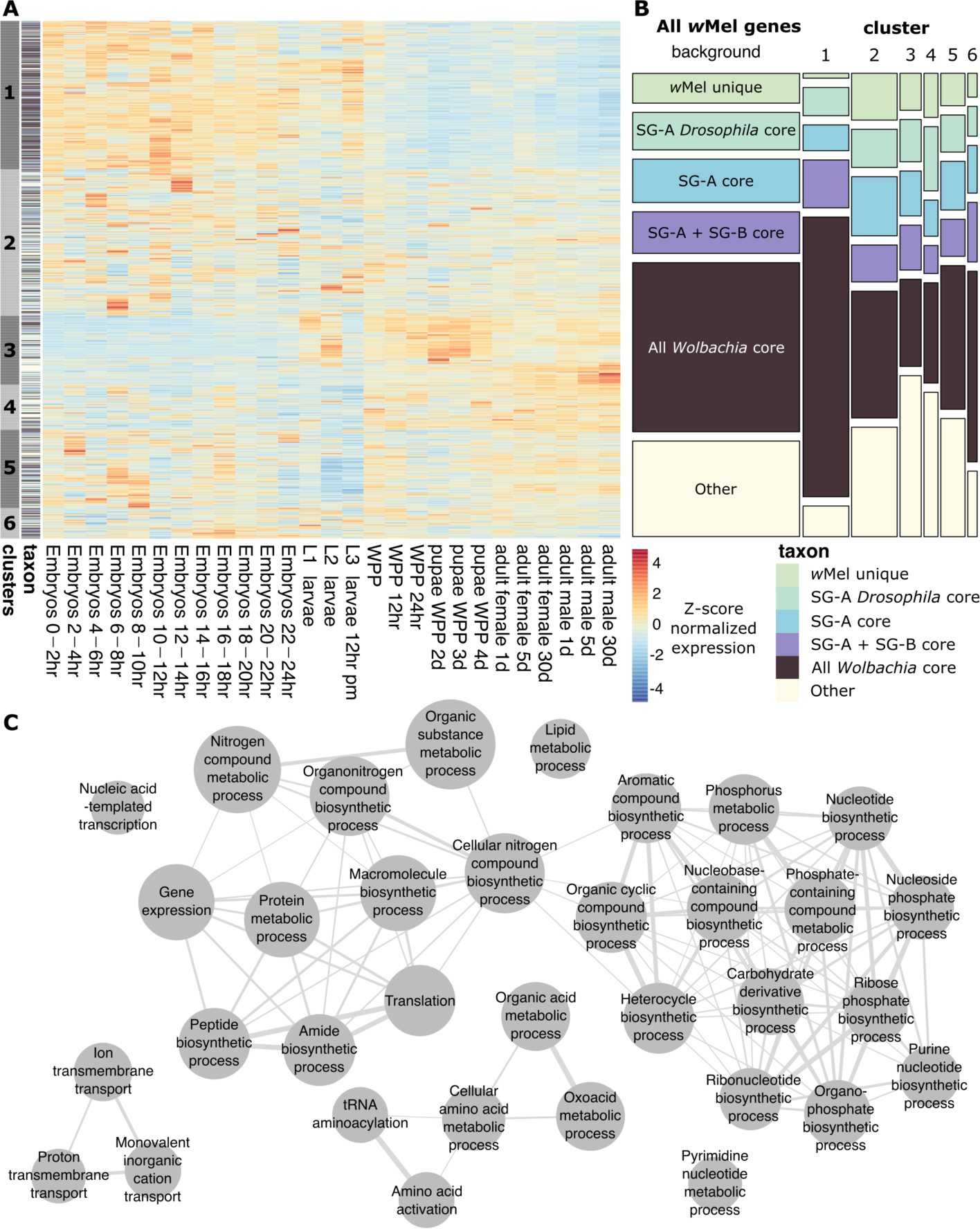
Global *Wolbachia* gene expression patterns during fly development. **(A)** Gene expression of all *w*Mel genes across fly development. *Wolbachia* transcriptomes were generated as part of the MODENCODE project (40, 41). Data are row normalized (z-score) and heatmap coloring indicates the relative expression of a given gene. Genes were clustered according to similarity of expression patterns across development, and each gene was assigned a taxonomic rank based on conservation relative to other *Wolbachia*. Taxonomic rank assignments include the following categories: (1) unique to *w*Mel, (2) core to the *Drosophila*-infecting clade within Supergroup A (*w*Mel, *w*Ri*, w*Au, *w*Rec, *w*Ha, *w*Suz), (3) core to Supergroup A, (4) core to Supergroups A and B, (5) core proteins across all *Wolbachia*, and, (6) all other *w*Mel proteins. **(B)** Mosaic plot showing the relative proportion of *w*Mel genes within a cluster at each taxonomic rank, relative to all expressed *w*Mel genes (“background”). Abbreviations: Supergroup (SG), white prepupae (WPP), larval instars 1, 2, and 3 (L1, L2, L3), 12 hours pre-metamorphosis (12 hr pm). **(C)** Network of significantly overrepresented GO terms (nodes) derived from cluster 1 genes. Edges indicate at least 20% shared genes, with thicker edges indicating more shared genes.

### Dynamics of *Wolbachia* gene expression and metabolism across fly development

Given the striking impact of *Wolbachia* infection on the larval transcriptome, and the historically strong focus on *Wolbachia* biology in adults, we leveraged an existing dataset to better assess (1) how *Wolbachia* might be interacting with juvenile hosts, and, if (2) patterns in *Wolbachia* gene expression might be functionally relevant to the changes we saw in larval gene expression. We identified several key features of *Wolbachia* gene expression across host development (Figure 2A): (1) there is a major shift in global *Wolbachia* gene expression that coincides with the transition to pupation (the transition between third instars (L3) and white pre-pupae (WPP)), (2) distinct clusters of genes have relatively high or low expression pre- and post-pupation, and, (3) genes with different evolutionary histories (e.g., core *Wolbachia* proteins, *w*Mel-specific genes) were non-randomly distributed across the expression clusters (Figure 2B; X-squared = 205.97, df = 25, p < 0.0001). For example, genes in cluster 1 were more highly expressed pre-pupation (Figure 2A), and significantly more likely to be a core *Wolbachia* gene (Figure 2B). In contrast, cluster 3 genes followed an opposite pattern: low expression pre-pupation and high expression post pupation. Additionally, the clustering algorithm identified two sub-clusters within cluster 3, and these largely correspond to genes that have high expression in pupae versus those with high expression in adult males. Cluster 3 was significantly overrepresented for transposition-related genes (Supplemental Table S9), and largely contained IS4/5 family transposases. In contrast, cluster 1 was significantly enriched for many GO terms related to core metabolism (Supplemental Table S8, Figure 2C). The strong upregulation of *Wolbachia*’s metabolic pathways in juvenile flies likely relates to *Wolbachia*’s own metabolic needs. However, this pattern might also be part of a regulatory strategy to limit negative impacts on the host and/or shift roles to better interface with different aspects of host biology across development. Regardless, the up regulation of *Wolbachia* metabolism in juveniles likely plays an important role in shifting the fly metabolic landscape. Finally, it is notable that the most evolutionarily conserved *Wolbachia* proteins are more highly expressed in juveniles, perhaps pointing more broadly towards the importance of these stages across other *Wolbachia*-host associations.

### *Wolbachia* is beneficial under nutrient limited conditions

Given the strong impact of *Wolbachia* on larval gene expression and the differential regulation of *Wolbachia* metabolic gene expression across fly development, we hypothesized that *Wolbachia* would impact fly developmental trajectories. To test this, we reared flies on 100% strength and 25% strength media and quantified pupation and adult emergence. We found a significant interaction between *Wolbachia*, media strength, and time that impacted fly pupation (F_1,1063_ = 24.7147, p < 0.0001) and adult emergence (F_1,1063_ = 22.2897, p < 0.0001). Additionally, we found a significant interaction between the presence of *Wolbachia* and the media strength (wild-type pupae: F_1,1063_ = 0.6517, p = 0.0063; wild-type adults: F_1,1063_ = 0.1746, p = 0.0062; *w*^145^ pupae: F_1,711_ = 8.8129, p = 0.0009; *w*^145^ adults: F_1,711_ = 19.0998, p < 0.0001), along with a significant impact of *Wolbachia* infection alone on fly pupation and adult emergence (wild-type pupae: F_1,1063_ = 0.2874, p < 0.0001; wild-type adults: F_1,1063_ = 0.5375, p < 0.0001; *w*^145^ pupae: F_1,711_ = 10.5483, p < 0.0001; *w*^145^ adults: F_1,711_ = 7.0718, p < 0.0001). For example, while the wild-type flies with and without *Wolbachia* reared on 100% strength media reached adulthood in 82% and 81% of cases, but on the 25% strength media these were reduced to 75% and 66%, respectively. The *Wolbachia*-mediated advantage was more prominent for a second fly genetic background, *w*^145^. Even on 100% strength media, the *Wolbachia*-free *w*^145^ flies experienced a 5% reduction in adult emergence relative to *Wolbachia*-infected flies (Figure 1D). When reared on 25% strength media, the *Wolbachia*-infected *w*^145^ flies were more than twice as likely to reach adulthood (70% versus 31%, Figure 1E). Across these assays, we found delays in entering pupation are approximately equal to the delay in adult emergence: *i.e.,* the time spent in metamorphosis did not change as a factor of *Wolbachia* infection. In summary, the *Wolbachia*-mediated advantage is driven by (1) *Wolbachia-*infected flies developing faster (1-5 days depending on genotype, Figures 1B and 1E) and a (2) a larger percentage of flies reaching adulthood.

Given that the effect size for the *Wolbachia* advantage was genotype dependent and that there was a more subtle impact of *Wolbachia* in the wild-type flies, we wondered if an even weaker media would result in a stronger advantage for the *Wolbachia*-infected flies. Conversely, *Wolbachia* could be a burden under more significant nutritional stress. Indeed, when flies were raised on 12.5% strength media, we again saw significant effects of the interaction between *Wolbachia* and time on the rate and success of pupation and adult emergence (pupae: F_1,347_ = 77.135, p < 0.0001; adults: F_1,347_ = 58.9296, p < 0.0001). However, *Wolbachia*-infected flies now developed slower than their uninfected counterparts (6-7 days delayed), but they pupated at higher rates (68% versus 61%) and were more likely to reach adulthood (54% versus 40%, Figure 1C). Furthermore, although *Wolbachia*-infected fly development was slower, it resulted in them attaining, on average, a 23% larger size at pupation (F_1,191_ = 32.076, p < 0.0001, Figure 1F). Additionally, *Wolbachia*-infected flies experienced lower levels of pupal mortality (35% versus 19%), perhaps related to the finding that flies that died during pupation were significantly smaller (F_1,191_ = 4.625, p = 0.0328, Figure 1F).

To determine if the developmental advantage was due to direct effects of *Wolbachia* or indirect effects via *Wolbachia*-mediated impacts on the gut microbiome, we performed the same developmental assay under axenic conditions (i.e., without a gut or food microbiome). Under axenic conditions, we saw a strong interactive effect of *Wolbachia* and time that resulted in significantly more, and faster, pupation and adult emergence for infected flies (Figure 1G; pupae: F_1,1920_ = 153.3796, p < 0.0001; adults: F_1,1920_ = 129.6080, p < 0.0001). Even on the 100% strength media, only 43% of *Wolbachia*-free flies pupated, compared to 61% of *Wolbachia*-infected flies. Again, *Wolbachia*-free flies had higher levels of pupal mortality: 21% of those that pupated did not emerge as adults, as compared to 13% of the *Wolbachia*-infected pupae. While removing the microbiota negatively impacted fly developmental timing as expected, this effect was exacerbated in the *Wolbachia*-free flies (Figures 1A and 1G). As compared to the conventionally-reared flies on 100% strength media (Figure 1A), axenic *Wolbachia*-infected flies experienced a two-day developmental delay, and *Wolbachia*-free flies were delayed three days (Figure 1G). The combination of axenic conditions and 25% strength media resulted in especially impaired fly development: less than 20% of flies reached adulthood (Figure 1H). While there were no significant differences in the proportion of flies that pupated or emerged as adults between *Wolbachia*-infected and uninfected flies in these high-mortality conditions (pupae: p = 0.9547; adults: p = 0.6210), it is notable that *Wolbachia*-infected flies on average started emerging five days ahead of the *Wolbachia*-free flies (Figure 1H). Finally, we measured the puparia of the axenic flies reared on 100% strength media (Figure 1I) and found that *Wolbachia* infection resulted in, on average, 10% larger pupal volumes (F_1,200_ = 11.69, p = 0.0008, Figure 1I), again indicating a significant developmental advantage.

### Pyrimidines contribute to the *Wolbachia* infection advantage

We reasoned that deficiencies in the diet supplemented by *Wolbachia* could be supplemented by exogenous nutrient addition to the fly food. Therefore, flies with and without *Wolbachia* would have the same growth parameters if this important nutrient was added. Specifically, we focused on the precursors (amino acids) and end-products (uridine and inosine) of *de novo* nucleotide synthesis, because of the related signatures in the fly and *Wolbachia* transcriptomes (Figures 1-2), and our previous data that purines play an important role in the symbiosis (20). We again prepared the 12.5% media which we then supplemented with these compounds. The developmental assays on supplemented media were repeated with the wild-type stocks, avoiding *w*^145^ because the *white* transporter is known to affect nucleotide metabolism (60). We again saw the *Wolbachia*-mediated developmental advantage in the 12.5% strength media (pupae: F_1,235_ = 13.9505, p = 0.0002; adults: F_1,235_ = 12.7568, p = 0.0004). Additionally, we saw a subtle but significant positive impact of *Wolbachia* in the 100% strength media as measured by adult emergence (pupae: F_1,235_ = 1.3002, p = 0.2538; adults: F_1,235_ = 28.0455, p < 0.0001), due to 6% pupal mortality in the *Wolbachia*-free flies that the *Wolbachia*-infected flies did not experience (Figure 4A). While the addition of casein benefited both *Wolbachia*-infected and uninfected flies, *Wolbachia*-free flies pupated at significantly increased rates relative to *Wolbachia*-infected flies (F_1,235_ = 9.8699, p = 0.0017). Again, pupal mortality was quite high in the *Wolbachia*-free flies (36% as compared to 9% with *Wolbachia*), resulting in significantly fewer numbers of adults (F_1,235_ = 6.9988, p = 0.0082). The addition of uridine (product of *de novo* pyrimidine synthesis) specifically benefited *Wolbachia*-free flies, but not the flies that had *Wolbachia* infections. Indeed, uridine supplementation resulted in a rescue phenotype where equivalent numbers of *Wolbachia-*infected and uninfected flies emerged as adults (p = 0.9281). Importantly, emergence of *Wolbachia*-free flies supplemented with uridine was not significantly different than the *Wolbachia*-infected flies on the standard 12.5% strength media (p = 0.9097) (Figure 4A). In fact, *Wolbachia*-free flies supplemented with uridine developed slightly faster than their infected counterparts (pupae: F_1,235_ = 2.2401, p = 0.1348; adults: F_1,235_ = 10.5842, p = 0.0011). The addition of inosine (product of *de novo* purine synthesis) was toxic and reduced pupation and emergence for all flies. As such there were no significant differences between *Wolbachia*-infected and uninfected flies in these high mortality (*i.e*., zero-inflated) conditions (pupae: F_1,235_ = 0.9217, p = 0.335; adults: F_1,235_ = 0.3519, p = 0.553). Simultaneously adding uridine and inosine resulted in more intermediate, albeit still quite lethal developmental phenotypes with no significant impact of *Wolbachia* (F_1,235_ = 1.3471, p = 0.2459; adults: F_1,235_ = 1.0400, p = 0.307).

Congruent with the previous developmental assays (Figure 3), *Wolbachia* infection had a significant impact on pupal size, both due to interacting effects with the media (F_5,470_ = 2.55, p = 0.0272), and *Wolbachia* alone (F_1,470_ = 37.31, p < 0.0001) (Figure 4B). Furthermore, *Wolbachia*-infected flies were significantly larger when reared on 100% (p = 0.0401), 12.5% (p = 0.0316), and casein-enriched 12.5% media (p < 0.0001). However, the addition of uridine to the 12.5% media resulted in similarly sized flies regardless of *Wolbachia* infection (p = 0.3685). In summary, *Wolbachia*-free flies were rescued to the same size and level of adult emergence as *Wolbachia*-infected counterparts in uridine (i.e., pyrimidine) supplemented media.

**Figure 3.**
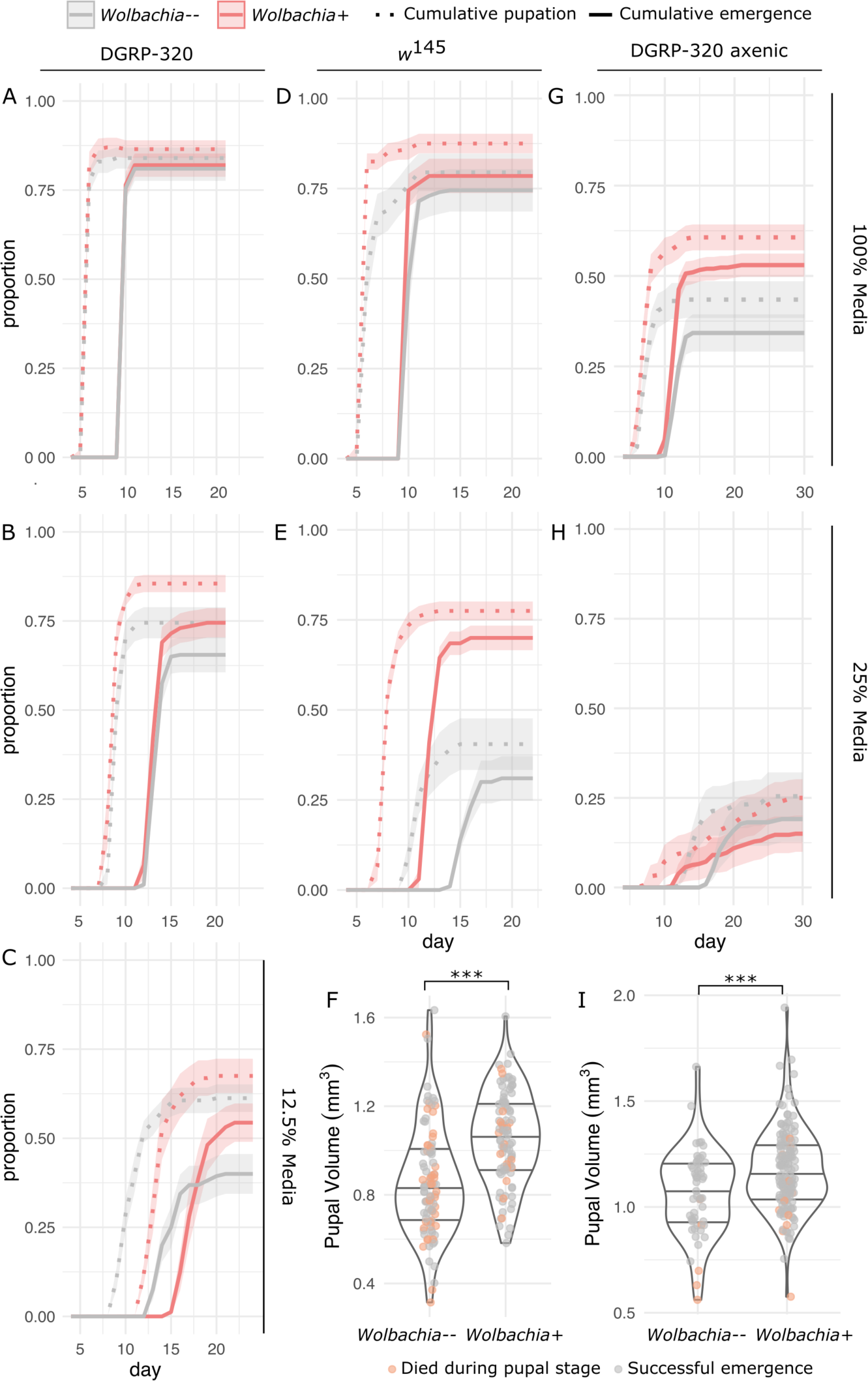
*Wolbachia* infection is beneficial under nutrient limited conditions. Flies (wild-type DGRP-320 and *w*^145^, with and without *Wolbachia*) were reared on different concentrations of media to test the impact of *Wolbachia* infection. Biological replicates included 20 larvae per each of 10 vials. Dotted and solid lines indicate cumulative pupation and adult emergence, respectively, with shaded regions defining standard error. All *Wolbachia*-infected treatments are in red, and *Wolbachia*-uninfected in grey. **(A)** Conventionally reared wild-type flies on 100%, **(B)** 25%, and **(C)** 12.5% strength media. **(D)** Conventionally reared *w*^145^ flies on 100% and **(E)** 25% strength media. **(F)** Pupal volumes of wild-type flies with and without *Wolbachia*, derived from the 12.5% media-reared flies in (C). **(G)** Axenic wild-type flies on 100% and **(H)** 25% strength media. **(I)** Pupal volumes of the axenic-reared flies on 100% strength media derived from (G). Orange datapoints (H-I) indicate pupae that did not eclose into adults. *** p < 0.001

We were curious about the fact that we also saw a size difference due to *Wolbachia* in the flies reared on 100% media (Figure 4B). Additionally, we noticed that the pupal sizes of the *Wolbachia*-infected flies followed a much more bimodal distribution than did the *Wolbachia* free-flies (Figure 4B). We wondered if this was due to differential impacts of *Wolbachia* on male versus female flies. To determine if sex was a contributing factor to the *Wolbachia*-mediated size difference, we measured sex-sorted wild-type pupae, reared on either 100% or 12.5% media, with and without *Wolbachia* (Figure 5). We found significant effects of sex and its interaction with *Wolbachia* or media on pupal size (sex\**Wolbachia*: F_1,194_ = 9.365, p = 0.0025; sex*media: F_1,194_ = 17.736, p < 0.0001; sex: F_1,194_ = 265.067, p < 0.0001). As in our previous assay, flies with *Wolbachia* produced significantly larger pupae than those without *Wolbachia* when reared on the 12.5% media (p < 0.0001). Additionally, this size increase with *Wolbachia* infection was more pronounced in females (41% larger than uninfected pupae) than in males (26% larger). In contrast to our previous assay, there were no significant differences in mean pupal size due to *Wolbachia* for either sex on 100% media (females: p = 0.9999; males: p = 0.8274). However, in these groups, there were significant differences in the distribution of pupal sizes. Specifically, while uninfected female pupae on 100% media followed a normal distribution (Shapiro-Wilk: W = 0.9778, p = 0.8513), the infected female pupal sizes significantly deviated from a normal distribution (Shapiro-Wilk: W = 0.9012, p = 0.0228). Similarly, pupal size distributions were significantly different between the *Wolbachia* infected and uninfected males derived from 100% media (Kolmogorov-Smirnov: D = 0.3896, p-value = 0.0267).

**Figure 4.**
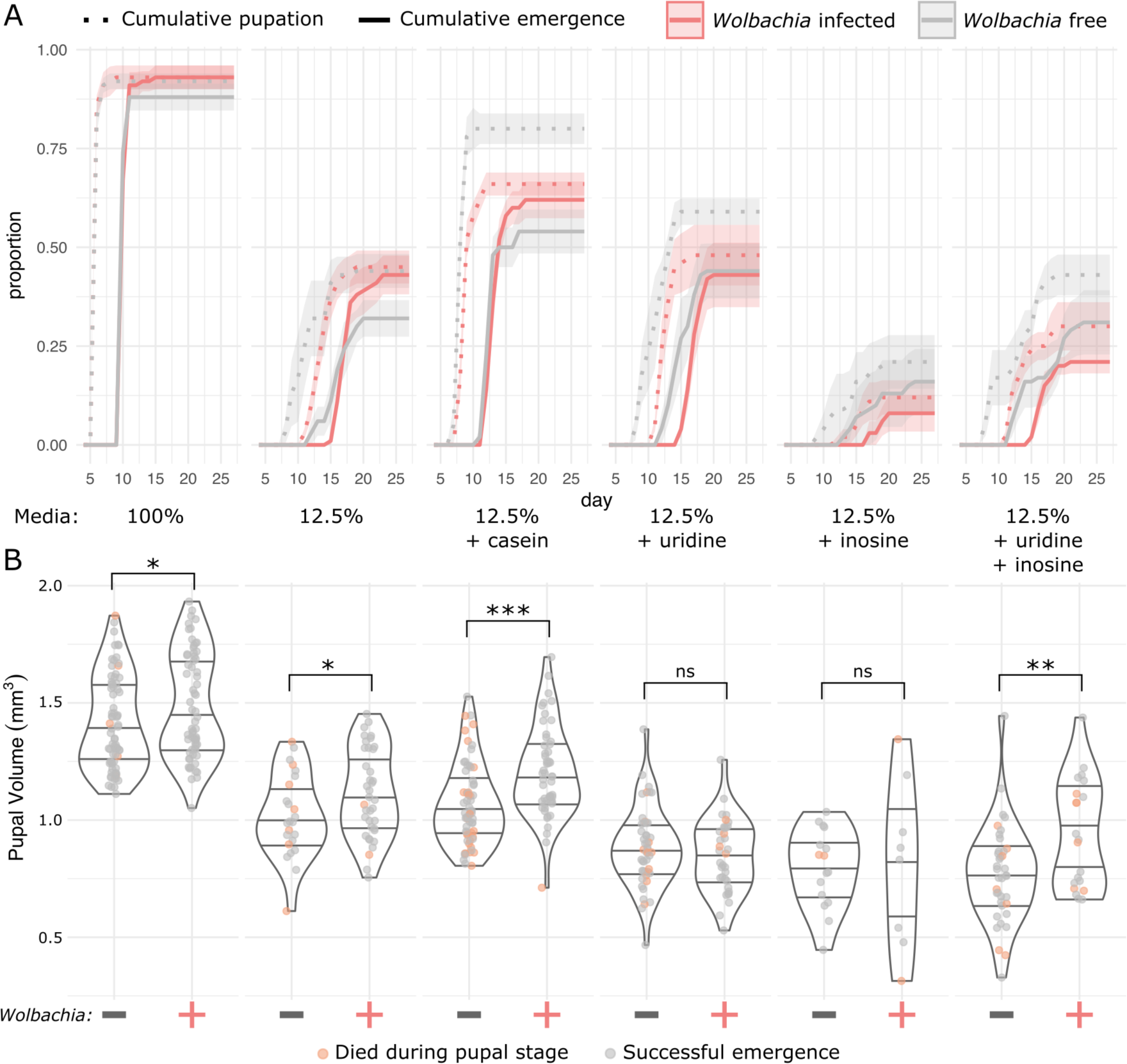
Uridine partially rescues *Wolbachia*-free flies. Wild-type flies were reared on 100% media, 12.5% media, or 12.5% media containing either casein, uridine, inosine, or both inosine and uridine. The media designations in the middle correspond to the figures above and below. Biological replicates included 20 larvae per each of five vials. **(A)** Dotted and solid lines indicate cumulative pupation and adult emergence, respectively, with shaded regions defining standard error. *Wolbachia*-infected treatments are in red and *Wolbachia*-uninfected are in grey. **(B)** Pupae from (A) were removed post-developmental assay, measured, and their volume was calculated. Orange datapoints indicate pupae that did not eclose into adults. Significance annotations for pupal sizes comparisons are the same regardless of whether flies that died during pupation were included. ***p < 0.001; **p < 0.01; *p < 0.05; ns = not significant.

**Figure 5.**
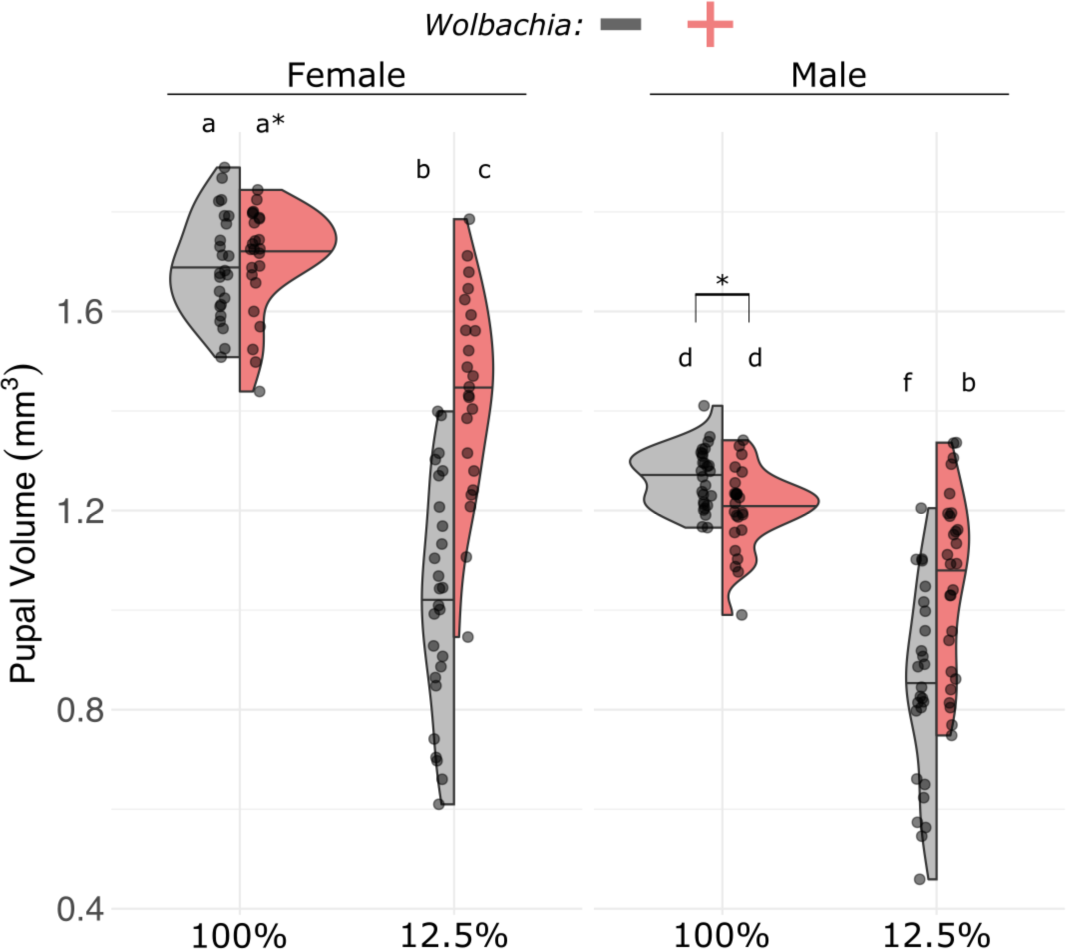
The *Wolbachia* size-advantage is more pronounced for female flies. Wild-type flies were reared on 100% or 12.5% media and collected during pupal stages P13-P14 (51). Pupae were sex sorted, measured, and their volume was calculated. Letters indicate statistically significant differences in mean pupal volume between groups (Tukey’s Honest Significant Difference test, p<0.05). Asterisks indicate significant differences in distributions.

### *Wolbachia* infection is responsive to fly pyrimidine biosynthesis gene expression

To determine whether *Wolbachia* infection was sensitive to changes in host metabolism, we knocked down individual genes within the fly’s *de novo* nucleotide synthesis pathways (Figure 6A) using *Wolbachia*-infected RNAi fly stocks (Figure 6B-C). Short-hairpin RNA (shRNA) expression was driven in the entire body of the fly by an Act5C-Gal4 driver. We compared gene expression and *Wolbachia* titer in knockdown flies to sibling controls (flies not expressing a gene-targeting shRNA, and instead with the TM6B balancer). We successfully knocked down four genes in the purine synthesis pathway (*prat2*, *ade2*, *ade5*, *CG11089*) and three genes in the pyrimidine synthesis pathway (*su(r)*, *Dhod*, *rudimentary-like*)(Figure 6B-C). Several gene knockdowns were lethal to fly development, including *rudimentary* in the de novo pyrimidine biosynthesis pathway and *CG2246*, *CG6767*, *Prat*, and *AdSL* in the de novo purine biosynthesis pathway. Broad-spectrum knockdown of *de novo* purine synthesis had no significant impacts on *Wolbachia* titer (gene*knockdown: F_3,31_ = 0.652, p = 0.588; gene: F_3,31_ = 0.644, p = 0.593; knockdown: F_1,31_ = 0.593, p = 0.447; Figure 6D). In contrast, knockdown of the *de novo* pyrimidine synthesis pathway resulted in significant increases in *Wolbachia* titer (F_1,38_ = 10.3916, p = 0.0026; Figure 6E). This pattern was specifically driven by the enzymatic loci (*Dhod*: p = 0.034 and *rudimentary-like* (*r-l*): p = 0.013). In contrast, knockdown of the negative regulator *sur(r)* had no effect on *Wolbachia* titer (p = 0.530), perhaps indicating that *Wolbachia* titers are responsive to low fly pyrimidine synthesis activity, but not elevated levels. To account for any effects of the TM6B balancer on measurements of *Wolbachia* abundance, we crossed the Act5C-Gal4 driver to a RNAi line expressing shRNA against a foreign gene: *Photinus pyralis* luciferase (UAS-anti-Ppyr\LUC). We found no significant changes in *Wolbachia* titer, or in nucleotide gene expression that could be driving the results in Figure 6B (Supplemental Figure S3). We then asked if the change in *Wolbachia* titer was tissue-specific by measuring the relative *Wolbachia* titers in ovaries and the remaining carcasses after knockdown of *r-l* (Figure 6F). There was no significant interaction of knockdown and tissue (F_1,32_ = 0.20, p = 0.658) or of tissue alone (F_1,32_ = 0.20, p = 0.658) on *Wolbachia* titers. We again saw a significant effect of *r-l* knockdown on *Wolbachia* titers (F_1,32_ = 13.07, p = 0.001) both in ovaries (p = 0.007) and in carcasses (p = 0.0127) (Figure 6F).

**Figure 6.**
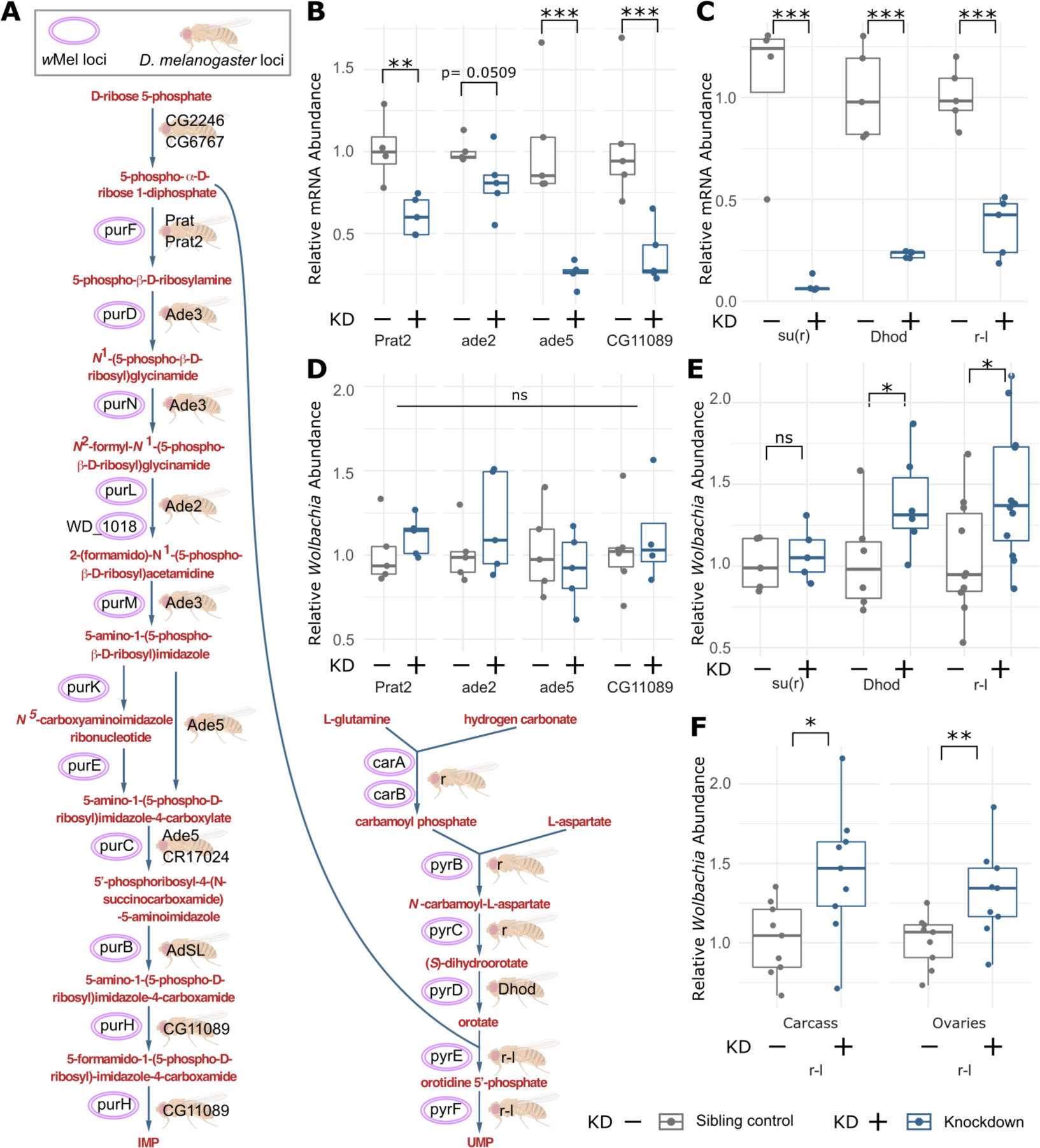
*Wolbachia* titers are responsive to fly pyrimidine synthesis gene expression. **(A)** Metabolic pathway for *de novo* nucleotide biosynthesis (purine: IMP, left; pyrimidine: UMP, right) in *Wolbachia* and *Drosophila melanogaster.* **(B)** Knockdown of fly purine biosynthetic genes was verified with qRT-PCR leveraging gene specific primers and normalization to 18S expression. **(C)** Knockdown of fly pyrimidine biosynthetic genes was verified with qRT-PCR leveraging gene specific primers and normalization to 18S expression. **(D)** Relative *Wolbachia* titers in whole adult females after purine gene knockdown. **(E)** Relative *Wolbachia* titers in whole adult females after pyrimidine knockdown. **(F)** Relative *Wolbachia* titers in female carcasses and ovaries after constitutive knockdown of *r-l*. In all cases, *Wolbachia* titers were quantified with qPCR using *Wolbachia*-specific primers and normalization to fly genome copy number via 18S amplification. All knockdowns leveraged short-hairpin RNA (shRNA) expression constitutively driven by Act5C-Gal4. Legend: *** p <0.001; ** p < 0.01; * p < 0.05; ns p = not significant.

### Interactive effects of *Wolbachia*, fly genotype, and diet impact fly metabolism

Considering the complex relationship between *Wolbachia* infection and *Drosophila* metabolism, we decided to further examine this relationship using semi-targeted LC-MS-based metabolomics to measure the relative abundance of 259 metabolites in two genetic strains (wild-type DGRP-320 and *w*^145^) raised on either 100% or 25% media (Figure 7; Supplemental Tables S10-12). The resulting data revealed that *Wolbachia* induced significant differences in the metabolome of both strains under both dietary conditions (Figure 7A-B). Moreover, the associated changes in the fly metabolome largely correlated with the strength of the developmental phenotype. For example, *Wolbachia* had reduced influence on the metabolome of wild-type flies grown on 100% media when compared with uninfected controls (Figure 7C,E), with only 11 metabolites exhibiting significant difference on the 100% food as compared to the 25 metabolites that were changed on the 25% media. Similarly, the total number of *Wolbachia-*associated metabolites that reached the cutoff threshold for significance in *w*^145^ was also lower in the 100% media (19 vs 29) (Figure 7D,F).

**Figure 7.**
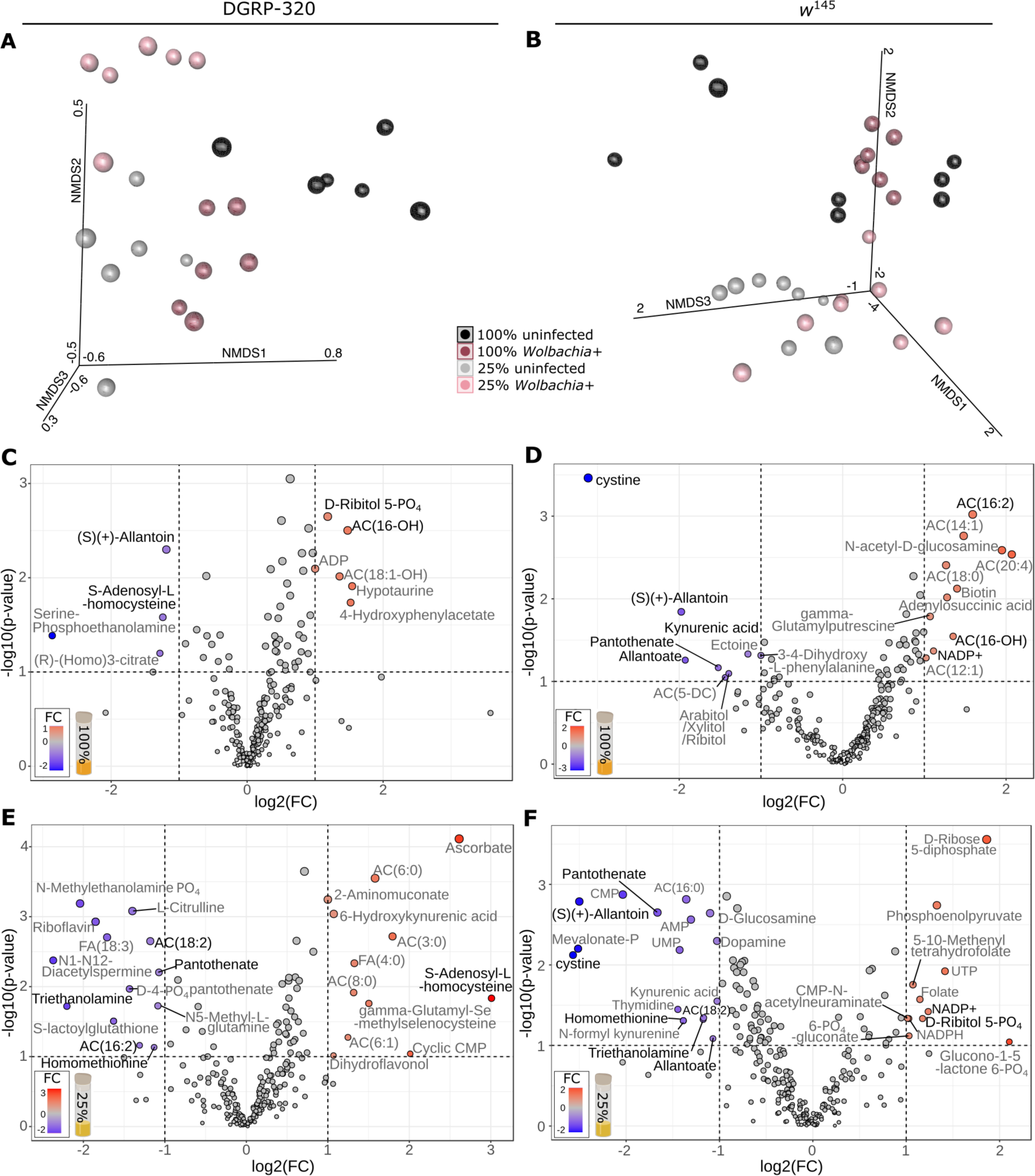
*Wolbachia* mediated impacts on the metabolome depend on host genotype and nutrition. **(A-B)** Three-dimensional non-metric multidimensional scaling (NMDS) plots indicating relative similarity of (A) DGRP-320 and (B) *w*^145^ samples. Point size indicates distance from the page, where larger points are closer. **(C-F)** Volcano plots indicating the relative abundance of specific metabolites in *Wolbachia* infected larvae relative to uninfected larvae: **(C)** DGRP-320 L2 larvae reared on 100% media. **(D)** *w*^145^ L2 larvae reared on 100% media. **(E)** DGRP-320 L2 larvae reared on 25% media. **(F)** *w*^145^ L2 larvae reared on 25% media. Metabolites with >2 log_2_ fold change (log2FC) and p<0.1 are indicated, and, for those that met these criteria in >2 conditions (e.g., panels) the metabolite name is in black font rather than grey. “Phosphate” is abbreviated with “PO_4_” to save space. “AC(#:#)” or “FA(#:#)” refers to acylcarnitines or fatty acids where numbers in parentheses indicate the total number of carbons in the chain, and the number of unsaturated bonds, respectively.

Beyond the total number of metabolites that were changed in *Wolbachia*-infected and uninfected larvae, several of the significantly altered compounds function within metabolic pathways that are clearly associated with *Wolbachia* infection. Specifically, in 3 of 4 analyses, *Wolbachia* infections results in reduced levels of the purine degradation products allantoin and allantoate (Figure 7C,D,F), whose production is associated with the antioxidant properties of uric acid (61). *Wolbachia* infected *w*^145^ flies also displayed decreased accumulation of kynurenic acid and cystine – amino acid derived molecules that are involved in the cellular response against oxidative stress (Figure 7D,F) (62–64). Our analysis also revealed decreases in pantothenate, a precursor to coenzyme A, thus suggesting that *Wolbachia* influences the abundance of a key molecule in mitochondrial metabolism. Finally, we would highlight the interactive impacts on S-adenosyl-L-homocysteine (SAH) abundance in the wild-type flies, an intermediate molecule in glutathione biosynthesis and thus directly involved in the cellular oxidative stress response.

## DISCUSSION

*Wolbachia* has long been appreciated for its diverse impacts on a range of ecdysozoan hosts. Increasingly, there is speculation as to the role *Wolbachia* plays in host metabolism more broadly, oftentimes in the context of specific metabolites (12, 20, 21). For example, *Wolbachia-*infection contributes to B-vitamin supplementation in a planthopper (19), bed bugs (15), and likely some species of solitary bees (65). Iron has been the subject of a number of investigations into *Wolbachia*-host interactions (16, 17, 66, 67). In *Drosophila melanogaster, Wolbachia* infection buffers against stresses associated with low or high concentrations of iron in the diet (67). *Wolbachia* infection is additionally associated with changes in host ferritin expression and iron homeostasis in other insects such as *Drosophila simulans* and the parasitoid wasp *Asobara tabida* (17). Given the range of metabolites and physiological processes that have been implicated in *Wolbachia*-host interactions, we focused initial experiments on using a more holistic approach to understanding fly development in the context of diet and *Wolbachia* infection.

Our transcriptomics results contained several surprising findings. First, a huge number of genes were differentially expressed due to *Wolbachia* infection: equivalent to 14% of the transcriptome. This number of DEGs is an order of magnitude larger than the number of DEGs identified in other analyses focused on adult transcriptomes (20). We were also surprised to see a clear impact on expression of the immune system. The prevailing hypothesis in the field has been that natively associated *Wolbachia* (e.g., *w*Mel for *Drosophila melanogaster*) do not trigger an immune response, whereas newly initiated infections (e.g., *w*Mel established in *Aedes* mosquitoes) do trigger an immune response (68). Indeed, while we did not see any induction of the immune system in *Wolbachia*-infected larvae, we did see strong down-regulation of many immunity-associated loci indicating some level of interaction between *Wolbachia* and host in this native association. Other notable findings in the transcriptomes include differential expression of major regulators of metabolism and development such as *Notch*, *InR*, and juvenile hormone signaling related proteins. Genes associated with the digestive system, fat body, muscles, and testes were also significantly more likely to be differentially expressed. While the first three tissues clearly point towards metabolic impacts, we found the pattern of testes-associated genes quite curious given that the analysis focused on L2s which had not been sex sorted. This pattern might reflect sex-specific differences in the timing of gonadal development: ovary morphogenesis does not begin until late L3, whereas testes begin to grow and differentiate substantially during embryogenesis (69). The earlier development of testes likely results in more power to detect *Wolbachia*-mediated impacts in these tissues as compared to in the developing ovaries, especially when using pooled, homogenized, whole animals. However, it is important to note that these metrics of tissue-biased expression are based on the MODENCODE dataset (37, 40, 41), which, as we note above, is derived from *Wolbachia*-infected flies. So, even given these caveats, there is clearly a *Wolbachia*-dependent difference in gene expression across a wide variety of processes.

Our gene expression data and understanding of *Wolbachia* genome evolution pointed us towards fly development and metabolism as an important regulator of the symbiosis. Indeed, we found that *Wolbachia* significantly impacts *Drosophila melanogaster* development, resulting in larger flies that were more likely to reach adulthood. The strength of this phenotype varied by fly genotype and the overall concentration of nutrition in the food: *Wolbachia* infection was highly beneficial under nutrient limited conditions. Removal of the extracellular microbial community indicated that this advantage was directly mediated by *Wolbachia*. In fact, *Wolbachia* significantly buffered flies against the stresses normally associated with removal of the microbiome (70, 71). Furthermore, *Wolbachia* titers were responsive to changes in the expression of fly pyrimidine synthesis, and the supplementation of pyrimidines rescued adult emergence of the *Wolbachia*-uninfected flies. Importantly, while the transcription of *Wolbachia*’s nucleotide metabolism pathways are quite high pre-pupation (Figure 2), the flies have comparatively low *Wolbachia* titers during that time (40). We propose that pyrimidines are a significant component of the developmental advantage provided by *Wolbachia*. During the ∼200-fold increase in larval body mass, there is a concomitant 25-fold increase in DNA and RNA content that creates a huge demand for pyrimidines, well exceeding the fly’s endogenous biosynthetic capacity (72). Considering that the rate of *Drosophila* development is highly sensitive to *de novo* pyrimidine synthesis (72), bacterially-derived nucleotides could give the host a significant advantage.

While we found that pyrimidines seemed to be important for the *Drosophila*-*Wolbachia* symbiosis, we did not see any changes in *Wolbachia* titer due to suppression of purine synthesis (Figure 6), nor did we see purine-mediated rescue of *Wolbachia*-uninfected flies (Figure 4). In fact, the addition of inosine to media negatively impacted flies. While altering the expression of fly purine metabolic gene expression did not impact the relative abundance of *Wolbachia* in our experiments, we cannot rule out the potential for other changes in *Wolbachia* gene expression or physiology. Nevertheless, these patterns are especially curious given that both metabolomics (Figure 7), and larval gene expression data (Figure 1) indicate significantly altered purine metabolism due to *Wolbachia*. And, we previously found an interactive effect of *Wolbachia* infection and the *de novo* purine biosynthesis pathway on virus replication (20), highlighting both the importance of purines to the symbiosis and the complexity of the metabolic landscape. Indeed, purine nucleotides act as signaling molecules that impact a range of functions. For example, activation of the adenosine receptor globally suppresses fly metabolism (73), which can cause insecticidal effects (74). Another possibility to explain the toxic effects of inosine supplementation (Figure 4) is misincorporation of inosine into RNAs. In mammalian cell lines, this can happen when inosine is in excess, ultimately impacting translation (75).

The contrast between purine and pyrimidine metabolism and their respective roles in the *Wolbachia*-host relationship led us to look more deeply into (1) how these pathways contrast with each other, and (2) how these pathways relate to other metabolic processes in the animal. Pyrimidines are synthesized by a series of metabolic reactions that use ribose sugar and the amino acids glutamine and aspartate to generate uridine-5’-monophosphate, UMP (76): the precursor for uracil, cytosine, and thymine. An enzyme in this pathway, dihydroorotate dehydrogenase (*Dhod*), is located in the inner mitochondrial membrane where it couples pyrimidine synthesis to electron transport (77, 78). This pathway is conserved in both *Drosophila* and *Wolbachia*, suggesting that infected flies generate pyrimidines in multiple compartments: the host cytoplasm/mitochondria and *Wolbachia*. Purine biosynthesis, again conserved between the fly and *Wolbachia*, is a comparatively more complicated pathway that also uses amino acids to build the nucleotide ring (Figure 6A). Additionally, an important early step in this pathway is the production of 5-phospho-α-D-ribose 1-diphosphate (PRPP) which is also used in pyrimidine biosynthesis (Figure 6A). In later steps, the enzyme AdSL generates 5-Aminoimidazole-4-carboxamide ribonucleotide (AICAR, the reactant for the next purine synthesis step, Figure 6A) and fumarate which is transported to the mitochondrion for the citric acid cycle. The connectivity of these two metabolic pathways, their links to amino acid and vitamin metabolism (e.g., folic acid derivatives), and the direct relationship with mitochondria implicates a much more complicated model than *Wolbachia* simply increasing the abundance of a singular limiting metabolite. For example *Wolbachia-*generated pyrimidines could reduce the biosynthetic burden normally imposed upon mitochondria – a model further supported by the ability of UMP to inhibit fly enzymes CAD and UMP synthase, which act as components of the fly’s pyrimidine biosynthetic pathway (79). Therefore, *Wolbachia*’s production of pyrimidines could simultaneously impact free amino acid pools (the nucleotide synthesis precursors), reduce mitochondrial burdens, and spatially segregate components of a normally inhibitory interaction.

Several other metabolites associated with mitochondrial metabolism and redox balance were significantly altered in *Wolbachia* infected animals. Decreases in purine catabolic products, kynurenic acid, cysteine, and SAH highlight metabolic pathways that are clearly associated with *Wolbachia* infection. *Wolbachia*-mediated changes in redox balance have been the subject of a number of previous investigations (80–85), though there are limited functional data to support the ultimate mechanisms responsible. A previously suggested model, based on metabolic differences in adult flies, hypothesized a reprogramming of mitochondrial metabolism, specifically towards non-oxidative metabolism combined with reduced insulin signaling and a hypoxia response (66). Our data largely support this hypothesis, and indeed, we do see downregulation of the insulin receptor and upregulation of a negative regulator in larvae with *Wolbachia*, plus impacts on redox balance. This same model and others (66, 86) further hypothesized that *Wolbachia* is exporting ATP (a purine), which contrasts with our findings that pyrimidines play a key role. However, given the link between these pathways, perhaps our supplementation of the highly limiting pyrimidines resulted in a reduced need for shunting PRPP to the pyrimidine pathway, ultimately allowing greater synthesis of IMP, which can later be converted to ATP. Whether *Wolbachia* are in fact exporting ATP remains to be empirically tested. Importantly, our understanding of *Wolbachia* physiology is largely limited to what we have inferred from genomic data and changes that we can detect in host physiology. While *Wolbachia* encode for an abundance of predicted transporters (3), we do not know if these proteins are functional, if their function agrees with bioinformatic predictions, or if specific metabolites are indeed transported in or out of *Wolbachia* cells.

Changes in the concentrations of the purine catabolic product SAH specifically point towards a model linking pathogen blocking with potential roles for *Wolbachia* in host metabolism. The observed changes in SAH are also quite intriguing considering that this molecule is both produced by S-adenosyl-L-methionine (SAM) methyltransferases and also inhibits the activity of these same enzymes (87). Since *Wolbachia* is proposed to enhance pathogen blocking by interacting with methylation of viral RNA molecules (88), our results suggest that this phenomenon could be the result of nutrient-dependent influences of *Wolbachia* on the host metabolism. The relative accumulation of SAH could be indicative of higher levels of methyltransferase activity. Notably, the interacting partner (*ipod*) of the fly’s methyltransferase (*dnmt2)* was strongly upregulated (68-fold) in the *Wolbachia*-infected larvae. *Wolbachia* may mediate effects on the host’s production of SAH, and/or *Wolbachia* might directly produce SAH, as the *Wolbachia* genome encodes for several putative methyltransferases (3). Indeed, our metabolomics approach does not allow us to identify which organism(s) any of the metabolite pools are derived from. However, given the overlap of several key metabolic pathways (Figure 6)(21) we hypothesize that many differences we detect are due to a combination of *Wolbachia*’s own metabolic activity and impacts on the host.

Our hypothesis that *Wolbachia*’s contributions to metabolism go beyond simple supplementation are further supported by our finding that *Wolbachia* decreases pupal mortality. This was especially notable during nutritional stress, suggesting that *Wolbachia*-free flies enter pupation despite insufficient metabolic reserves. This effect was most pronounced when flies were reared on an excess of protein (Figure 4). Importantly, amino acids are a key nutritional cue for determining the timing of pupation (i.e., signaling critical weight) due to direct and indirect effects on the timing and intensity of ecdysone pulses (89–92). An influx of amino acids without other essential nutrients needed for growth might impact nutritional signaling and thus the timing of critical weight, perhaps triggering flies to pupate early without sufficient energy stores needed for metamorphosis. *Wolbachia*’s import of amino acids may reduce the concentrations of free amino acids in these protein enriched conditions, which could delay signaling critical weight, allowing flies to accumulate more metabolic reserves before pupation.

The role of *Wolbachia* as a nutritional symbiont raises numerous questions about the cell biology of *Wolbachia* and how the relationship with the host is regulated. Indeed, the patterns of *Wolbachia* gene expression do seem to be attuned to fly development (Figure 2) which could be a strategy for supporting the exponential growth that occurs during larval stages and mitigating fitness impacts of the infection during reproductive stages. Is *Wolbachia* specifically providing pyrimidines or other key metabolites (*e.g.,* via nucleotide transporters)? Or are flies farming and consuming their *Wolbachia* infections (*e.g.,* via autophagy) and pyrimidines so happen to be abundant in *Wolbachia* cells and also a limiting metabolite for the fly? As is the case for many symbioses, it is not always clear if a change in the symbiont (*e.g.,* gene expression or titer), is due to the microbe detecting and responding to the host’s status, or the host releasing control of the microbe and “allowing” a change in microbial physiology. *Wolbachia* do encode for a number of proteins that would allow them to detect their environment and regulate gene expression accordingly (93). However, each *Wolbachia* is enclosed by a host-derived vesicle (94, 95), so perhaps the host tightly regulates *Wolbachia*’s “experience”. Changes in *Wolbachia* titer and physiology across development and under different nutritional scenarios likely reflect these evolutionary balancing acts, as is evident by the complex manner by which rapamycin feeding and dietary amino acid supplementation influence *Wolbachia* titers (96).

While beneficial nutritional symbioses in insects are not rare, most of the systems for which this is described are of a much more binary nature. Insects that exclusively feed on unbalanced diets (e.g., blood, plant sap) have obligate microbial symbionts to synthesize metabolites that the host cannot generate *de novo*: the partners complement each other (97, 98). Here, both flies and *Wolbachia* have their own complete pathways for nucleotide biosynthesis, which perhaps has led us to overlook the relevance of this biosynthetic redundancy. Our finding that *Wolbachia* significantly impacts the developmental timing, size, and susceptibility to nutritional stress in a well-studied model such as *Drosophila melanogaster* raises the question of the broader relevance of *Wolbachia* infections to insect metabolism and development.

## DECLARATIONS

### Acknowledgements

Research reported in this publication was supported by the National Institute of General Medical Sciences of the National Institutes of Health under award number R35GM150991 and the National Institute of Allergy and Infectious Disease of the National Institutes of Health under award number R21AI175957, both to ARIL. ILGN is supported by the National Institute of Allergy and Infectious Diseases of the National Institutes of Health under R01AI144430. JMT is supported by the National Institute of General Medical Sciences of the National Institutes of Health under R35GM119557. The authors thank Richard Hardy for support and feedback. Research was further made possible by Flybase (NIH 5U41HG000739), and stocks obtained from the Bloomington *Drosophila* Stock Center (NIH P40OD018537).

## Data Availability

Raw RNA-seq data have been deposited in the NCBI Short Read Archive under BioProject PRJNA1111126.

Supplemental Table S1. Fly stocks

Supplemental Table S2. Primers

Supplemental Table S3. Expression data for all expressed genes in the *Drosophila melanogaster* L2 dataset.

Supplemental Table S4. Differentially expressed *Drosophila melanogaster* L2 genes.

Supplemental Table S5. Statistical results for overrepresented gene sets (PANGEA). Supplemental Table S6. Statistical results for overrepresented terms and clusters (STRING).

Supplemental Table S7. Taxon and cluster assignments for *w*Mel loci.

Supplemental Table S8. *w*Mel cluster 1 statistical results for overrepresented biological process GO terms.

Supplemental Table S9. *w*Mel cluster 3 statistical results for overrepresented biological process GO terms.

Supplemental Table S10. Metabolomics metadata

Supplemental Table S11. DGRP-320 metabolite abundance data Supplemental Table S12. *w*145 metabolite abundance data

Supplemental Figure S1. Full sized, all-labeled version of RNA-seq network (main text Figure 1C).

Supplemental Figure S2. Pupal measurement.

Supplemental Figure S3. Knockdown controls.

## REFERENCES

1. R. Kaur et al., Living in the endosymbiotic world of *Wolbachia*: A centennial review. Cell Host & Microbe 29(6), 879–893(2021).

2. O. Duron et al., The diversity of reproductive parasites among arthropods: *Wolbachia* do not walk alone. BMC Biol. 6, 1–12(2008).

3. M. Wu et al., Phylogenomics of the reproductive parasite *Wolbachia* pipientis *w*Mel: A streamlined genome overrun by mobile genetic elements. PLoS Biol. 2, 327–341(2004).

4. J. H. Werren, W. Zhang, L. R. Guo, Evolution and phylogeny of *Wolbachia* -reproductive parasites of arthropods. Proc. R. Soc. Lond. B 261, 55–63(1995).

5. N. Bensaadi-Merchermek, J.-C. Salvado, C. Cagnon, S. Karama, C. Mouchès, Characterization of the unlinked 16S rDNA and 23S-5S rRNA operon of *Wolbachia pipientis*, a prokaryotic parasite of insect gonads. Gene 165, 81–86(1995).

6. R. Zug, P. Hammerstein, Still a host of hosts for *Wolbachia*: Analysis of recent data suggests that 40% of terrestrial arthropod species are infected. PLoS One 7 (2012).

7. K. Hilgenboecker, P. Hammerstein, P. Schlattmann, A. Telschow, J. H. Werren, How many species are infected with *Wolbachia*? -A statistical analysis of current data. FEMS Microbiol. Lett. 281, 215–220(2008).

8. F. Rousset, D. Bouchon, B. Pintureau, P. Juchault, M. Solignac, *Wolbachia* endosymbionts responsible for various alterations of sexuality in arthropods. Proc. R. Soc. Lond. B 250, 91–98(1992).

9. K. Fenn, M. Blaxter, *Wolbachia* genomes: revealing the biology of parasitism and mutualism. Trends Parasitol. 22, 60–65(2006).

10. R. Stouthamer, J. A. J. Breeuwer, G. D. D. Hurst, *Wolbachia pipientis*: Microbial manipulator of arthropod reproduction. Annu. Rev. Microbiol. 53, 71–102(1999).

11. C. A. Hamm et al., *Wolbachia* do not live by reproductive manipulation alone: infection polymorphism in *Drosophila suzukii* and *D. subpulchrella*. Mol. Ecol. 23, 4871–4885(2014).

12. R. Zug, P. Hammerstein, Bad guys turned nice? A critical assessment of Wolbachia mutualisms in arthropod hosts. Biological Reviews 90, 89–111(2015).

13. A. R. I. Lindsey, T. Bhattacharya, I. L. G. Newton, R. W. Hardy, Conflict in the intracellular lives of endosymbionts and viruses: A mechanistic look at *Wolbachia*-mediated pathogen-blocking. Viruses 10, 141 (2018).

14. E. P. Caragata, H. L. Dutra, P. H. Sucupira, A. G. Ferreira, L. A. Moreira, *Wolbachia* as translational science: controlling mosquito-borne pathogens. Trends Parasitol. 37, 1050–1067(2021).

15. T. Hosokawa, R. Koga, Y. Kikuchi, X. Y. Meng, T. Fukatsu, *Wolbachia* as a bacteriocyte-associated nutritional mutualist. Proc. Natl. Acad. Sci. 107, 769–774(2010).

16. A. C. Gill, A. C. Darby, B. L. Makepeace, Iron necessity: the secret of *Wolbachia*’s success? PLoS Negl. Trop. Dis. 8, e3224 (2014).

17. N. Kremer, et al., *Wolbachia* interferes with ferritin expression and iron metabolism in insects. PLoS Path. 5, e1000630 (2009).

18. V. Geoghegan et al., Perturbed cholesterol and vesicular trafficking associated with dengue blocking in *Wolbachia*-infected *Aedes aegypti* cells. Nat. Commun. 8, 526 (2017).

19. J.-F. Ju et al., *Wolbachia* supplement biotin and riboflavin to enhance reproduction in planthoppers. The ISME journal 14, 676–687(2020).

20. A. R. Lindsey, T. Bhattacharya, R. W. Hardy, I. L. Newton, *Wolbachia* and virus alter the host transcriptome at the interface of nucleotide metabolism pathways. mBio 12 (2021).

21. I. L. Newton, D. W. Rice, The Jekyll and Hyde symbiont: could *Wolbachia* be a nutritional mutualist? J. Bacteriol. 202 (2020).

22. L. W. Beukeboom, Size matters in insects–an introduction. Entomol. Exp. Appl. 166, 2–3(2018).

23. C. Dmitriew, L. Rowe, The effects of larval nutrition on reproductive performance in a food-limited adult environment. PLoS One 6, e17399 (2011).

24. B. Taborsky, The influence of juvenile and adult environments on life-history trajectories. Proc. R. Soc. Lond. B 273, 741–750(2006).

25. J. H. Werren, D. M. Windsor, *Wolbachia* infection frequencies in insects: Evidence of a global equilibrium? Proc. R. Soc. Lond. B 267, 1277–1285(2000).

26. W. Huang et al., Natural variation in genome architecture among 205 *Drosophila melanogaster* Genetic Reference Panel lines. Genome Res. 24, 1193–1208(2014).

27. M. W. Jones, L. C. Fricke, C. J. Thorpe, L. O. V. Esch, A. R. I. Lindsey, Infection dynamics of cotransmitted reproductive symbionts are mediated by sex, tissue, and development. Appl. Environ. Microbiol. 88, e00529–00522(2022).

28. G. dos Santos et al., FlyBase: introduction of the *Drosophila melanogaster* Release 6 reference genome assembly and large-scale migration of genome annotations. Nucleic Acids Res. 43, D690–D697(2014).

29. B. Li, C. N. Dewey, RSEM: accurate transcript quantification from RNA-Seq data with or without a reference genome. BMC Bioinformatics 12, 323 (2011).

30. A. Dobin et al., STAR: ultrafast universal RNA-seq aligner. Bioinformatics 29, 15–21(2013).

31. R Core Team (2014) R: A language and environment for statistical computing. (URL http://www.R-project.org/, R Foundation for Statistical Computing, Vienna, Austria).

32. C. Soneson, M. I. Love, M. D. Robinson, Differential analyses for RNA-seq: transcript-level estimates improve gene-level inferences. F1000Research 4 (2015).

33. M. D. Robinson, D. J. McCarthy, G. K. Smyth, edgeR: a Bioconductor package for differential expression analysis of digital gene expression data. Bioinformatics 26, 139–140(2010).

34. D. J. McCarthy, Y. Chen, G. K. Smyth, Differential expression analysis of multifactor RNA-Seq experiments with respect to biological variation. Nucleic Acids Res. 40, 4288–4297(2012).

35. A. Yim et al., mitoXplorer, a visual data mining platform to systematically analyze and visualize mitochondrial expression dynamics and mutations. Nucleic Acids Res. 48, 605–632(2020).

36. F. Marchiano, M. Haering, B. H. Habermann, The mitoXplorer 2.0 update: integrating and interpreting mitochondrial expression dynamics within a cellular context. Nucleic Acids Res. 50, W490–W499(2022).

37. Y. Hu et al., PANGEA: a new gene set enrichment tool for *Drosophila* and common research organisms. Nucleic Acids Res. 51, W419–W426(2023).

38. N. T. Doncheva, J. H. Morris, J. Gorodkin, L. J. Jensen, Cytoscape stringApp: Network analysis and visualization of proteomics data. Journal of Proteome Research (2018).

39. P. Shannon et al., Cytoscape: a software environment for integrated models of biomolecular interaction networks. Genome Res. 13, 2498–2504(2003).

40. F. Gutzwiller et al., Dynamics of *Wolbachia pipientis* gene expression across the *Drosophila melanogaster* life cycle. G3: Genes| Genomes| Genetics 5, 2843–2856(2015).

41. B. R. Graveley et al., The developmental transcriptome of *Drosophila melanogaster*. Nature 471, 473–479(2011).

42. M. Chung et al., A meta-analysis of *Wolbachia* transcriptomics reveals a stage-specific *Wolbachia* transcriptional response shared across different hosts. G3: Genes, Genomes, Genetics 10, 3243–3260 (2020).

43. L. C. Fricke, A. R. Lindsey, Identification of parthenogenesis-inducing effector proteins in *Wolbachia*. Genome Biol. Evol., evae036 (2024).

44. M. Lechner et al., Proteinortho: detection of (co-) orthologs in large-scale analysis. BMC Bioinformatics 12, 1–9(2011).

45. R. Kolde, M. R. Kolde, Package ‘pheatmap’. R package 1, 790 (2015).

46. S. X. Ge, D. Jung, R. Yao, ShinyGO: a graphical gene-set enrichment tool for animals and plants. Bioinformatics 36, 2628–2629(2020).

47. S. Bainbridge, M. Bownes, Staging the metamorphosis of *Drosophila melanogaster*. Journal of Embryology and Experimental Morphology 66, 57–80(1981).

48. T. Reis, Effects of synthetic diets enriched in specific nutrients on *Drosophila* development, body fat, and lifespan. PLoS One 11, e0146758 (2016).

49. F. Demontis, N. Perrimon, Integration of Insulin receptor/Foxo signaling and dMyc activity during muscle growth regulates body size in Drosophila. (2009).

50. L. Moss-Taylor, A. Upadhyay, X. Pan, M.-J. Kim, M. B. O’Connor, Body size and tissue-scaling is regulated by motoneuron-derived activinß in *Drosophila melanogaster*. Genetics 213, 1447–1464(2019).

51. S. P. Bainbridge, M. Bownes, Staging the metamorphosis of Drosophila melanogaster. Development 66, 57–80(1981).

52. K. J. Livak, T. D. Schmittgen, Analysis of relative gene expression data using real-time quantitative PCR and the 2− ΔΔCT method. methods 25, 402–408(2001).

53. J. E. Cox, C. S. Thummel, J. M. Tennessen, Metabolomic studies in *Drosophila*. Genetics 206, 1169–1185(2017).

54. H. Li, J. M. Tennessen, Preparation of *Drosophila* larval samples for gas chromatography-mass spectrometry (GC-MS)-based metabolomics. JoVE (Journal of Visualized Experiments), e57847 (2018).

55. T. Nemkov, J. A. Reisz, S. Gehrke, K. C. Hansen, A. D’Alessandro, High-throughput metabolomics: isocratic and gradient mass spectrometry-based methods. High-Throughput Metabolomics: Methods and Protocols, 13–26(2019).

56. D. Bates, M. Maechler, B. Bolker, S. Walker, lme4: Linear mixed-effects models using Eigen and S4. R package version 1 (2014).

57. J. Oksanen et al., vegan: Community Ecology Package. R package version 2.0–1. http://CRAN.R-project.org/package=vegan (2015).

58. J. Xia, N. Psychogios, N. Young, D. S. Wishart, MetaboAnalyst: a web server for metabolomic data analysis and interpretation. Nucleic Acids Res. 37, W652–W660(2009).

59. T. Bhattacharya, I. L. G. Newton, R. W. Hardy, Wolbachia elevates host methyltransferase expression to block an RNA virus early during infection. PLoS Path. 13, e1006427 (2017).

60. A. Sasaki, T. Nishimura, T. Takano, S. Naito, S. K. Yoo, *white* regulates proliferative homeostasis of intestinal stem cells during ageing in *Drosophila*. Nature Metabolism 3, 546–557(2021).

61. A. J. Hilliker, B. Duyf, D. Evans, J. P. Phillips, Urate-null rosy mutants of *Drosophila melanogaster* are hypersensitive to oxygen stress. Proc. Natl. Acad. Sci. 89, 4343–4347(1992).

62. M. Conrad, H. Sato, The oxidative stress-inducible cystine/glutamate antiporter, system xc−: cystine supplier and beyond. Amino Acids 42, 231–246(2012).

63. R. Lugo-Huitrón et al., On the antioxidant properties of kynurenic acid: free radical scavenging activity and inhibition of oxidative stress. Neurotoxicol. Teratol. 33, 538–547(2011).

64. K. Afshar, P. Gönczy, S. DiNardo, S. A. Wasserman, *fumble* encodes a pantothenate kinase homolog required for proper mitosis and meiosis in *Drosophila melanogaster*. Genetics 157, 1267–1276(2001).

65. M. Gerth, C. Bleidorn, Comparative genomics provides a timeframe for *Wolbachia* evolution and exposes a recent biotin synthesis operon transfer. *Nat*. Micro. 2, 16241 (2016).

66. D. Currin-Ross et al., The metabolic response to infection with *Wolbachia* implicates the insulin/insulin-like-growth factor and hypoxia signaling pathways in *Drosophila melanogaster*. Frontiers in Ecology and Evolution 9, 623561 (2021).

67. J. C. Brownlie, et al., Evidence for metabolic provisioning by a common invertebrate endosymbiont, *Wolbachia pipientis*, during periods of nutritional stress. PLoS Path. 5, e1000368 (2009).

68. E. Rancès, Y. H. Ye, M. Woolfit, E. A. McGraw, S. L. O’Neill, The relative importance of innate immune priming in *Wolbachia*-mediated dengue interference. PLoS Path. 8, e1002548 (2012).

69. N. Camara, C. Whitworth, M. Van Doren, The creation of sexual dimorphism in the *Drosophila* soma. Current Topics in Developmental Biology 83, 65–107(2008).

70. N. A. Broderick, B. Lemaitre, Gut-associated microbes of *Drosophila melanogaster*. Gut Microbes 3, 307–321(2012).

71. D. N. Lesperance, N. A. Broderick, Microbiomes as modulators of *Drosophila melanogaster* homeostasis and disease. Current Opinion in Insect Science 39, 84–90(2020).

72. D. Falk, R. Bellamy, M. el Kouni, N. F, A genetic and dietary study of the physiology of pyrimidine synthesis in *Drosophila melanogaster*. J. Insect Physiol. 26, 735–740(1980).

73. A. Bajgar, T. Dolezal, Extracellular adenosine modulates host-pathogen interactions through regulation of systemic metabolism during immune response in Drosophila. PLoS Path. 14, e1007022 (2018).

74. M. Fang, Y. Chai, G. Chen, H. Wang, B. Huang, N6-(2-hydroxyethyl)-adenosine exhibits insecticidal activity against Plutella xylostella via adenosine receptors. PLoS One 11, e0162859 (2016).

75. J. H. Schroader et al., Disease-associated inosine misincorporation into RNA hinders translation. Nucleic Acids Res. 50, 9306–9318(2022).

76. D. R. Evans, H. I. Guy, Mammalian pyrimidine biosynthesis: fresh insights into an ancient pathway. J. Biol. Chem. 279, 33035–33038(2004).

77. J.-J. Chen, M. E. Jones, The cellular location of dihydroorotate dehydrogenase: relation to de novo biosynthesis of pyrimidines. Archives of Biochemistry and Biophysics 176, 82–90(1976).

78. J. M. Rawls, Genetic complementation and enzyme correlates at the locus encoding the last two steps of de novo pyrimidine biosynthesis in *Drosophila melanogaster*. Molecular and General Genetics MGG 184, 174–179(1981).

79. M. Löffler, E. A. Carrey, W. Knecht, The pathway to pyrimidines: The essential focus on dihydroorotate dehydrogenase, the mitochondrial enzyme coupled to the respiratory chain. Nucleosides Nucleotides Nucl. Acids 39, 1281–1305(2020).

80. L. Brennan, J. Haukedal, J. Earle, B. Keddie, H. Harris, Disruption of redox homeostasis leads to oxidative DNA damage in spermatocytes of *Wolbachia*-infected *Drosophila simulans*. Insect Mol. Biol. 21, 510–520(2012).

81. R. Zug, P. Hammerstein, *Wolbachia* and the insect immune system: what reactive oxygen species can tell us about the mechanisms of *Wolbachia*–host interactions. Frontiers in Microbiology 6, 1201 (2015).

82. X. Pan et al., *Wolbachia* induces reactive oxygen species (ROS)-dependent activation of the Toll pathway to control dengue virus in the mosquito *Aedes aegypti*. Proc. Natl. Acad. Sci. 109, E23–E31(2012).

83. D. Monnin et al., Influence of oxidative homeostasis on bacterial density and cost of infection in *Drosophila*–*Wolbachia* symbioses. J. Evol. Biol. 29, 1211–1222(2016).

84. Z. S. Wong, J. C. Brownlie, K. N. Johnson, Oxidative stress correlates with *Wolbachia*-mediated antiviral protection in *Wolbachia*-*Drosophila* associations. Appl. Environ. Microbiol. 81, 3001–3005(2015).

85. L. J. Brennan, B. A. Keddie, H. R. Braig, H. L. Harris, The endosymbiont *Wolbachia pipientis* induces the expression of host antioxidant proteins in an *Aedes albopictus* cell line. PLoS One 3 (2008).

86. A. C. Darby et al., Analysis of gene expression from the *Wolbachia* genome of a filarial nematode supports both metabolic and defensive roles within the symbiosis. Genome Res. 22, 2467–2477(2012).

87. R. Kumar, R. Srivastava, R. K. Singh, A. Surolia, D. N. Rao, Activation and inhibition of DNA methyltransferases by S-adenosyl-L-homocysteine analogues. Bioorganic & Medicinal Chemistry 16, 2276–2285(2008).

88. T. Bhattacharya, et al., Differential viral RNA methylation contributes to pathogen blocking in *Wolbachia*-colonized arthropods. PLoS Path. 18, e1010393 (2022).

89. S. Layalle, N. Arquier, P. Léopold, The TOR pathway couples nutrition and developmental timing in *Drosophila*. Dev. Cell 15, 568–577(2008).

90. C. Géminard, E. J. Rulifson, P. Léopold, Remote control of insulin secretion by fat cells in Drosophila. Cell Metabolism 10, 199–207(2009).

91. J. Colombani et al., A nutrient sensor mechanism controls *Drosophila* growth. Cell 114, 739–749(2003).

92. M. J. Texada, T. Koyama, K. Rewitz, Regulation of body size and growth control. Genetics 216, 269–313(2020).

93. A. R. I. Lindsey, Sensing, Signaling, and Secretion: A review and analysis of systems for regulating host interaction in *Wolbachia*. Genes 11, 813 (2020).

94. K.-O. Cho, G.-W. Kim, O.-K. Lee, *Wolbachia* bacteria reside in host Golgi-related vesicles whose position is regulated by polarity proteins. PLoS One 6, e22703 (2011).

95. P. M. White et al., Reliance of *Wolbachia* on high rates of host proteolysis revealed by a genome-wide RNAi screen of *Drosophila* cells. Genetics 205, 1473–1488(2017).

96. L. R. Serbus, et al., The impact of host diet on *Wolbachia* titer in *Drosophila*. PLoS Path. 11, e1004777 (2015).

97. N. A. Moran, G. R. Plague, J. P. Sandström, J. L. Wilcox, A genomic perspective on nutrient provisioning by bacterial symbionts of insects. Proc. Natl. Acad. Sci. 100, 14543–14548(2003).

98. N. A. Moran, Symbiosis. Curr. Biol. 16, R866–871(2006).

