## Supplemental Figures for "The intracellular symbiont *Wolbachia* alters *Drosophila* development and metabolism to buffer against nutritional stress"

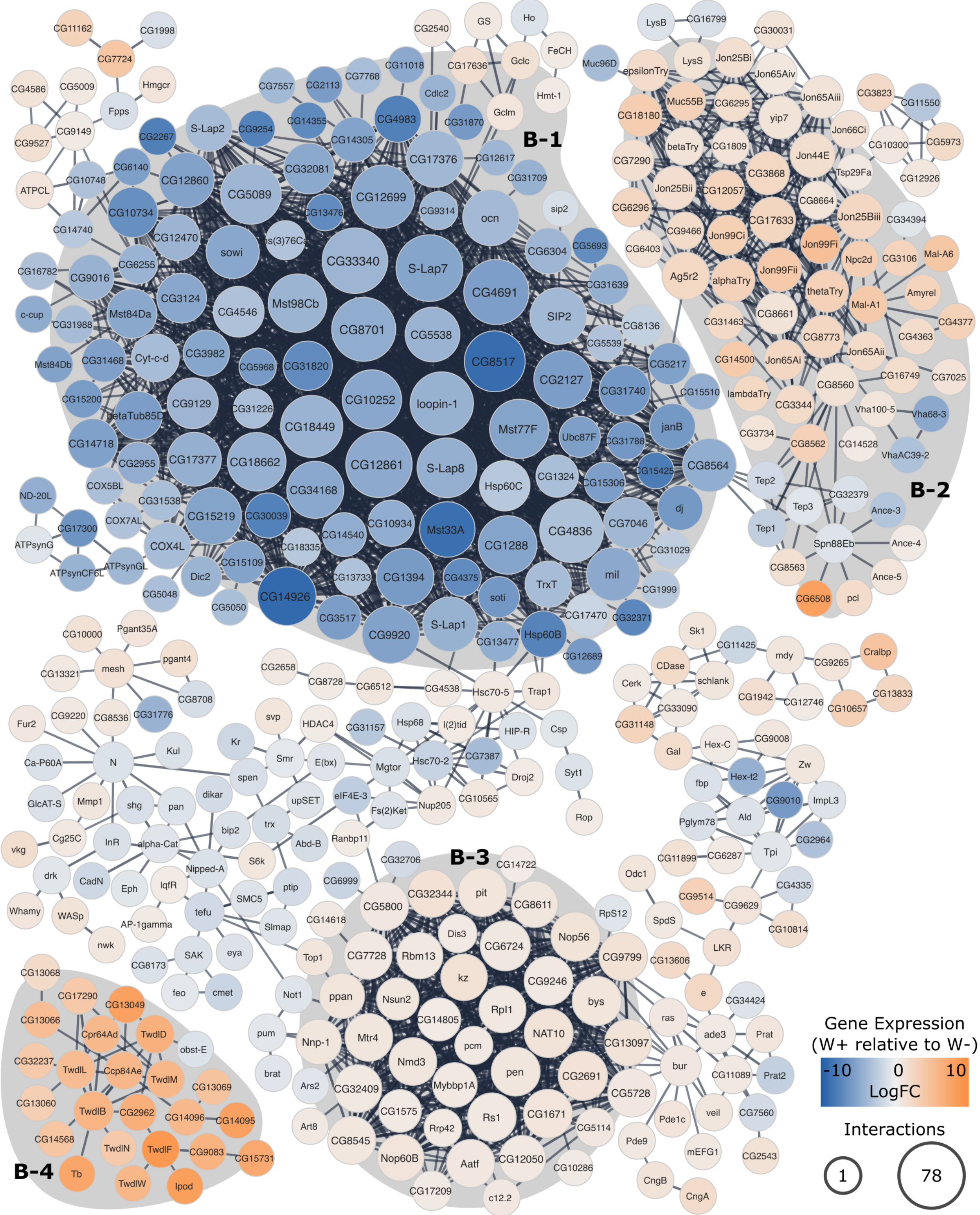


**Figure S1. Full sized, all-labeled version of RNA-seq network (Figure 1B).** Core protein-protein interaction networks within the differentially expressed gene set. Nodes are colored according to their change in gene expression relative to uninfected larvae (with orange indicating higher expression in *Wolbachia* infected larvae). The size of each node corresponds to the number of interactions that protein has with other nodes in the network. Shaded regions are clusters of the gene expression network that are significantly overrepresented with the following functional terms: **(B-1)** uncharacterized peptidases, transmembrane transport, protein localization to microtubule, **(B-2)** signal, serine proteases, extracellular, hydrolase, digestion, carboxypeptidase, transmembrane transport, **(B-3)** ribosome biogenesis and tRNA modification, and **(B-4)** signal, cuticle development, chitin.

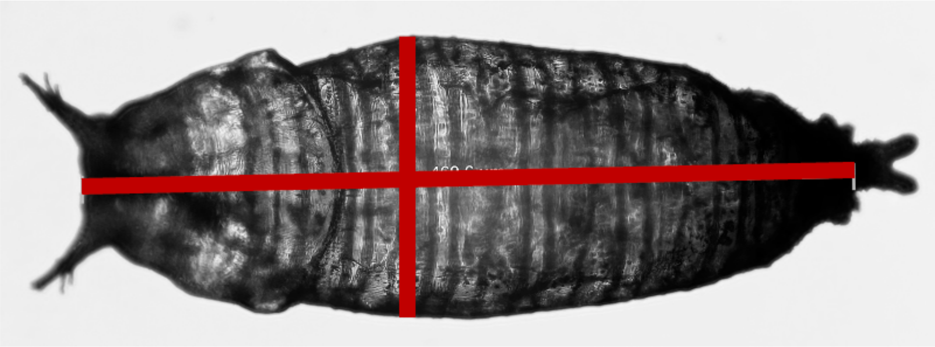

**Figure S2. Pupal measurements.** Pupal volume was calculated based on length and width (red lines) and assuming a prolate spheroid shape [$V = (4/3)\,\pi\,(\text{width}/2)^2\,(\text{length}/2)$].

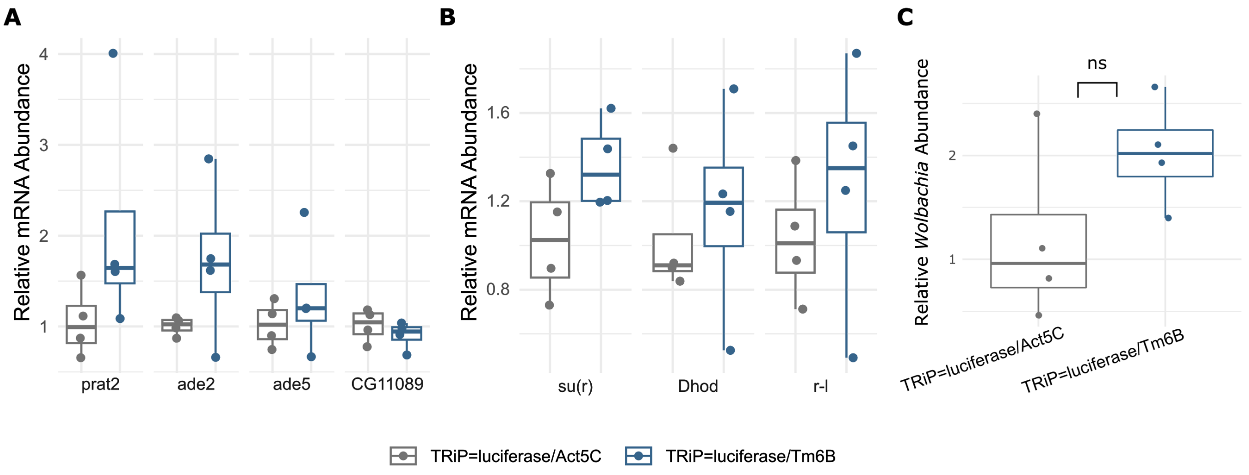


**Figure S3. Validation of knockdown controls.** Impact of balancer chromosomes on (A) purine gene expression, (B) pyrimidine gene expression, and (C) *Wolbachia* titer. **(A)** The TM6B balancer had no significant impacts on purine gene expression (gene*balancer: $F_{3,24} = 1.311$, p = 0.2939; gene: $F_{3,24} = 1.311$, p = 0.2939; knockdown: $F_{1,24} = 4.146$, p = 0.0529). Importantly, while the mean expression of the purine loci may have been marginally higher in the presence of the balancer chromosome, this means that we would underestimate the effects of knockdown due to an even larger difference between the sibling and knockdown flies (see main text). **(B)** The TM6B balancer had no significant impacts on pyrimidine gene expression (gene*balancer: $F_{2,18} = 0.227$, p = 0.799; gene: $F_{2,18} = 0.227$, p = 0.799; knockdown: $F_{1,18} = 1.180$, p = 0.292). **(C)** There was no significant difference in *Wolbachia* titer due to the balancer chromosome (t-test, t = 1.8404, df = 8, p = 0.1153). Again, the trend towards marginally higher *Wolbachia* titer in the presence of the balancer only means that we would underestimate the effects of knockdown.
